# The Long Noncoding RNA FEDORA is a Cell-Type- and Sex-Specific Regulator of Depression

**DOI:** 10.1101/2021.11.30.470628

**Authors:** Orna Issler, Yentl Y van der Zee, Aarthi Ramakrishnan, Sunhui Xia, Alexander K Zinsmaier, Chunfeng Tan, Wei Li, Caleb J Browne, Deena M Walker, Marine Salery, Angélica Torres-Berrío, Rita Futamura, Julia E Duffy, Benoit Labonte, Carol A Tamminga, Jeffrey L Dupree, Yan Dong, James W Murrough, Li Shen, Eric J Nestler

## Abstract

Women suffer from depression at twice the rate of men, but the underlying molecular mechanisms are poorly understood. Here, we identify dramatic baseline sex differences in expression of long noncoding RNAs (lncRNAs) in human postmortem brain tissue that are profoundly lost in depression. One such lncRNA, RP11-298D21.1 (which we termed FEDORA), is enriched in oligodendrocytes and neurons and upregulated in several cortical regions of depressed females but not males. We found that virally-expressing FEDORA selectively either in neurons or in oligodendrocytes of prefrontal cortex promoted depression-like behavioral abnormalities in female mice only, changes associated with cell-type-specific regulation of synaptic properties, myelin thickness, and gene expression. We also found that blood FEDORA levels have diagnostic significance for depressed women. These findings demonstrate the important role played by lncRNAs, and FEDORA in particular, in shaping the sex-specific landscape of the brain and contributing to sex differences in depression.

## Introduction

Major depressive disorder (MDD) is a devastating psychiatric syndrome and among the leading causes of disability worldwide ^1, 2^. Available treatments, including antidepressant medications and psychotherapy, have limited response and remission rates, with more than half of patients remaining at least partly treatment-resistant ^3, 4^. There are pronounced sex differences in MDD, as women are twice as likely to suffer from depression as men ^1, 5^. Moreover, women tend to have a different subset of symptoms which are often more severe ^5^, and respond differentially to mechanistically divergent antidepressants compared to men ^6^. These sex differences are partially attributable to a higher sensitivity of women’s stress responses ^7^ and to fluctuations in ovarian hormones ^8, 9^. However, these mechanisms do not explain why only a subset of women develop depression ^10^. Shedding light on the molecular processes associated with sex differences in MDD could promote the development of sex-specific psychiatric clinical tools.

Epigenetic processes have been proposed as important mediators of gene-environment interactions that lead to MDD ^3, 11^. One important arm of epigenetic regulation are long non-coding RNAs (lncRNAs), a class of transcripts longer than 200 nucleotides with no apparent protein-coding role ^12, 13^. Notably, about forty percent of lncRNAs are brain-specific, with many of these molecules arising in primate evolution and coinciding with an increase in brain size ^14^. LncRNAs have similar structural features to mRNAs, however, they are often expressed at low levels and exhibit striking region- and cell-type-specific patterns ^15^. Functionally, lncRNAs can act as scaffolds, guides, or decoys, leading to changes in expression of protein-coding genes ^16^. We and others reported regulation of specific lncRNAs in postmortem brain tissue of MDD patients ^17, 18^ and rodent stress models ^19, 20^. However, a global analysis of baseline sex differences in lncRNA expression levels in the brain, and how this pattern is altered in MDD, has not yet been performed.

The present study reports robust baseline sex differences in lncRNA expression patterns in several limbic brain regions examined postmortem from healthy control (HC) subjects that are dramatically lost in MDD. We highlight one such human-specific lncRNA, RP11-298D21.1, which we named FEDORA, that is expressed to a lower degree in female HCs compared to males across several brain regions examined. This pattern is reversed in MDD: FEDORA is upregulated in females with MDD compared to HCs, with no change seen in males. FEDORA expression in brain predominates in oligodendrocytes and to a lesser extent in neurons. We hypothesis that this lncRNA is a key sex-specific and cell-type-specific regulator of mood. To test this hypothesis, we virally-expressed FEDORA in prefrontal cortex (PFC) selectively in either neurons or oligodendrocytes in mice of both sexes, followed by comprehensive phenotyping. We report that, while FEDORA expression in both cell types promotes depression-like behavior in female mice only, this effect is mediated by opposing transcriptional changes in neurons compared to oligodendrocytes. We also show that expression of FEDORA in PFC neurons alters their electrophysiological activity, while expression of FEDORA in oligodendrocytes regulates myelin thickness. Finally, we show that FEDORA levels in circulating blood are upregulated only in females with MDD, with normalization of FEDORA expression correlating with the degree of clinical improvement seen in response to the rapidly-acting antidepressant ketamine. Together, this study highlights the importance of lncRNAs in the brain, both in maintaining baseline sex differences and in MDD pathophysiology, and suggests that FEDORA may serve as a key regulator of these processes.

## Results

### Abnormal Expression of FEDORA in Female MDD

In a recent study, we reported robust sex- and brain-site-specific regulation of lncRNAs in postmortem brain tissue of subjects with MDD compared to HCs ^17^. Utilizing this rich dataset, the present work focused on lncRNAs that demonstrate baseline sex differences, that is, are differentially expressed between male vs female HC subjects (Fold change > 30% and p < 0.05). Comparing MDD with HC subjects, we noted a dramatic loss of these baseline sex differences in lncRNA expression. In the ventral medial PFC (vmPFC), we identified 762 lncRNAs with baseline sex differences, with 65.9% of these lncRNAs losing their baseline pattern of sex-specific expression in MDD subjects (Fig. 1a). We found a similar phenomenon in each of the five other brain regions examined, which included dorsolateral PFC (dlPFC), orbitofrontal cortex (OFC), ventral subiculum (vSUB), anterior insula (aINS), and nucleus accumbens (NAc) (Extended Data Fig. 1a-h), and this effect was more pronounced when compared to the same analysis performed on protein-coding genes (PCGs; e.g., vmPFC = 45.1%; Extended Data Fig. 1). We observed two common patterns of regulation: the first was lncRNAs exhibiting lower expression in female vs male HCs that are upregulated in female MDD patients but downregulated (or not affected) in male MDD patients (Fig. 1c), and the second where lncRNAs exhibiting higher expression in females vs males HCs are downregulated in female MDD patients but upregulated (or not affected) in male MDD patients (Fig. 1d). These findings suggest that lncRNAs play a key role in regulating sex differences in the brain and that this pattern is corrupted by MDD.

**Fig. 1.**
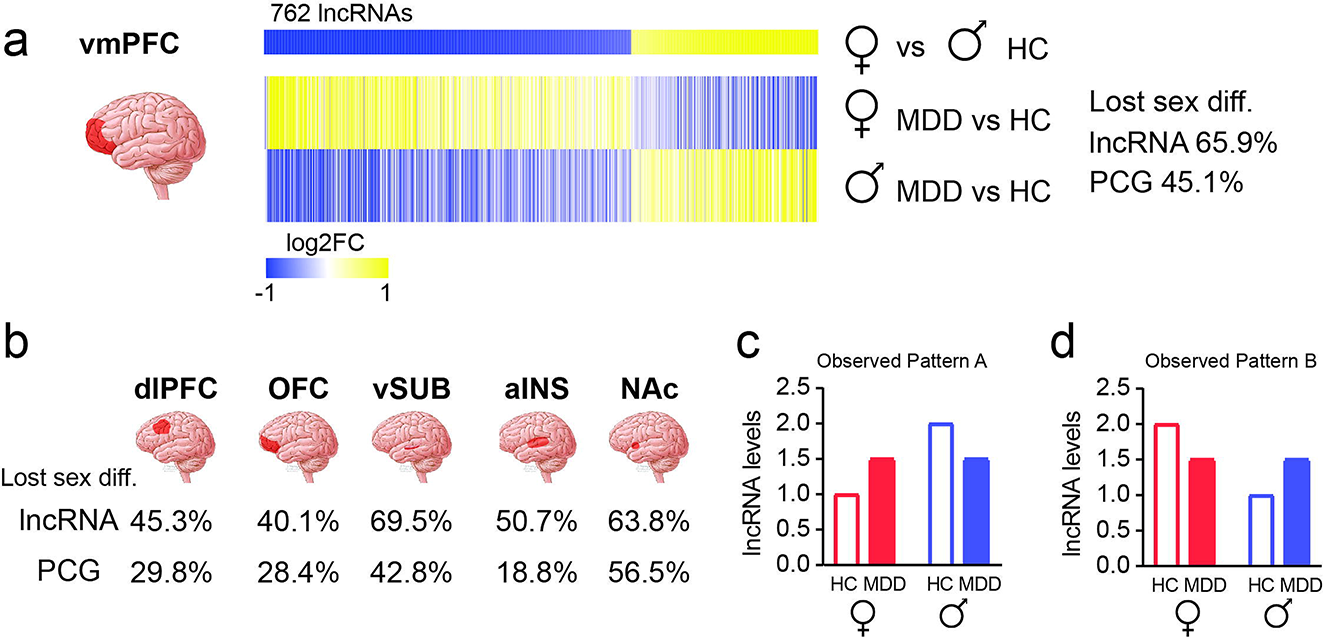
Baseline sex differences in lncRNA expression are lost in MDD. **(a)** Heatmaps demonstrating that lncRNAs in ventral medial prefrontal cortex (vmPFC) that exhibit baseline sex differences between female and male HCs (differential expression Fold change>30% and P< 0.05), are robustly oppositely altered in females vs males with MDD compared to HCs. Yellow represents upregulation and blue downregulation between log_2_fold change=±1. (**b**) Baseline sex differences are also lost in MDD in the dorsal lateral prefrontal cortex (dlPFC), orbitofrontal cortex (OFC), ventral subiculum (vSUB), anterior insula (aINS), and nucleus accumbens (NAc). The percentage of lncRNAs and PCGs that lose this baseline sex difference in MDD is indicated. **(c-d)** Bar graphs representing the most commonly observed patterns of regulation of lncRNAs that lose baseline sex differences in MDD. RNA-seq dataset is from ^26^. n=9-13 per group. MDD = major depressive disorder; HC = healthy control.

We hypothesize that lncRNAs that lose their baseline sex differences in MDD may have a critical mechanistic role in the molecular processes leading to sex differences in the pathophysiology of MDD. To identify such specific candidates, we probed the list of lncRNAs that demonstrated baseline sex differences (422 lncRNAs on average across brain regions) and filtered for those that broadly lost their baseline sex differences in MDD (73 lncRNAs in 4 or more of the brain regions analyzed). Next, we selected for lncRNAs that show a significant correlation in their expression levels to at least one PCG within our datasets (R> 0.587 or R< −0.595; 19 lncRNAs), are abundant in the brain (CPM>1; 15 lncRNAs), and are differentially expressed in MDD in either sex (Fold change>30% and P<0.05). This analysis revealed an antisense lncRNA called RP11-298D21.1 or AC009063.2 (Fig. 2a). This lncRNA displays a baseline sex difference, as it is expressed at lower levels in female vs male HCs, and is upregulated in MDD females compared to HCs across all brain regions studied, with no abnormal expression seen in MDD males (Fig. 2b). Based on this expression pattern, we named the lncRNA FEmale DepressiOn lncRNA (FEDORA). We validated these results for the vmPFC using a combined cohort including the same samples used for RNA-seq plus an additional cohort. We noted that FEDORA is upregulated in females with MDD compared to HCs (Two-way ANOVA, F interaction_(1,75)_=5.556, P=0.021, post hoc Fisher LSD, HC:female vs MDD:female t_(75)_=1.946, P=0.0409). FEDORA is also expressed at higher levels in male compared to female HCs, consistent with a reversal of this baseline sex difference in MDD (post hoc Fisher LSD HC:female vs HC:male t_(75)_=2.269, P=0.0261) (Fig. 2c).

**Fig. 2.**
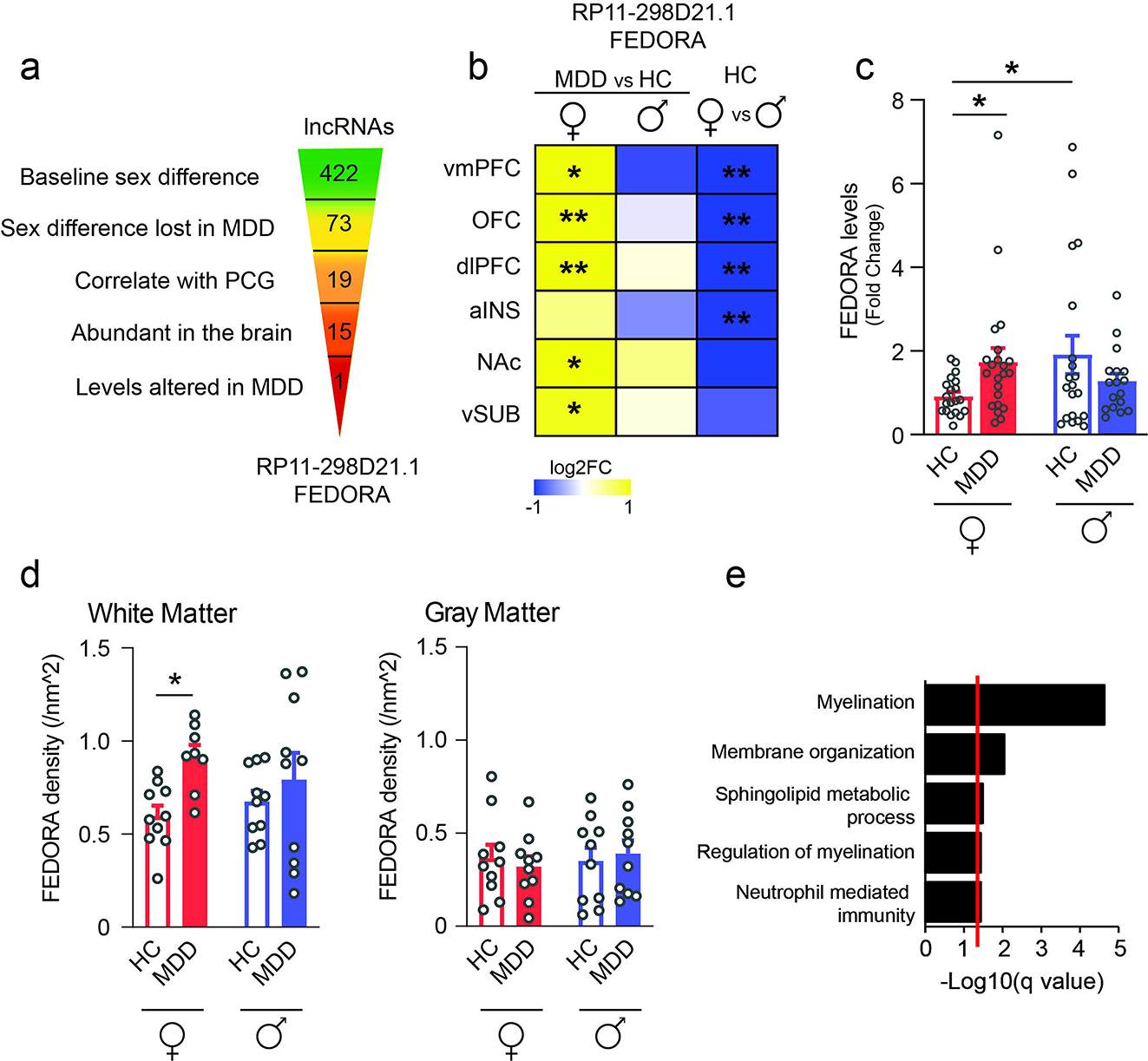
FEDORA lncRNA, a human oligodendrocyte- and neuronal-enriched transcript, is upregulated in female MDD. **(a)** Schematic representation of lncRNA filtering process identifying RP11-298D21.1 (FEDORA). LncRNAs with baseline sex differences (averaged across brain regions) were refined to those that: 1) lost this baseline difference in MDD in at least 4 brain regions, 2) exhibit significant correlation with at least one protein-coding gene (PCG) within our female postmortem brain dataset, 3) are abundant in the human brain (CMP>1), and 4) are differentially expressed in MDD (FC>30% and P<0.05). **(b)** Heatmaps representing RNA-seq results of the differential expression of FEDORA in female, but not male, MDD compared to HCs, as well as a baseline sex difference in HCs between female and male. * p<0.05; **p<0.01. n=9-13 per group. **(c)** Bar graph of qPCR validation results from a combined cohort of the samples used for RNA-seq plus additional samples showing higher levels of FEDORA in males compared to females in the HCs as well as upregulation of FEDORA in females with MDD, with no effect of MDD seen in males. * p<0.05 N=17-21 per group. Bars represent mean ± SEM and dots represent individual data points. **(d)** Bar graph of quantification of FISH for FEDORA in rostral anterior cingulate cortex (rACC) of an independent cohort of human brain postmortem samples. The data show that FEDORA is expressed at higher levels in white matter than in gray matter and is upregulated in females with MDD compared to HCs. * p<0.05 N=10 per group. **(e)** Bar graph representing top enriched GO ontology terms of PCGs (n=360) whose expression levels are significantly correlated with FEDORA in the female brain. The red line represents q=0.05.

FEDORA is a human-specific antisense RNA of *Cdh13*, which encodes cadherin-13 ^21^. FEDORA is enriched in brain ^22^ (Extended Data Fig. 2a), where it is predominantly expressed in oligodendrocytes and in neurons to a lesser extent ^23^ (Extended Data Fig. 2b). We validated the regulation of FEDORA in MDD and its cell-type-specific expression pattern using in situ hybridization in the rostral anterior cingulate cortex (rACC) from an independent cohort of MDD and HC subjects of both sexes. We found that FEDORA co-localizes with markers for both neurons and oligodendrocytes (Extended Data Fig. 2c-d), with higher levels of FEDORA in white matter than in gray matter (Two-way ANOVA in control subject, F main effect tissue _(1,36)_=9.621, P=0.0037) (Fig. 2d). Within white matter, we found higher levels of FEDORA in females with MDD compared to control, but not in males (Two-way ANOVA, F main effect group _(1,34)_=5.733, P=0.0223) (Fig. 2d). Gene ontology analysis performed on the list of 360 PCGs that significantly correlate with FEDORA levels in the female brain within our dataset implicated myelination-related processes (Fig. 2e), supporting a potential role for FEDORA in oligodendrocyte function.

### Behavioral Effects of FEDORA Expression in Mouse mPFC

To determine whether FEDORA has a causal sex- and cell-type-specific role in promoting MDD, we “humanized” mice for this transcript by virally expressing it selectively in either neurons or oligodendrocytes. First, we developed a neuron-specific tool to express FEDORA in mouse mPFC along with a GFP reporter (Neuro-FEDORA), or GFP only as control (Neuro-GFP), using Herpes simplex virus (HSV), which is highly neurotropic (Fig. 3a). We validated that this tool promotes FEDORA expression using qPCR (Extended Data Fig. 3a,b) and that the expression of FEDORA is neuron-specific by co-localizing Neuro-GFP with neuronal, but not with oligodendrocyte or astrocytic, markers using immunohistochemistry (Extended Data Fig. 3c). Next, we infected the mPFC of female and male adult mice with Neuro-FEDORA or Neuro-GFP, and subsequently performed a battery of tests to measure anxiety- and depression-related behaviors. In female mice, Neuro-FEDORA compared to Neuro-GFP led to a longer latency to feed in the novelty suppressed feeding test (Student’s *t*-test, *t*_15_=2.180, P=0.0456) (Fig. 3b), more marbles buried in the marble burying test (Student’s *t*-test, *t*_15_=2.351, P=0.0326) (Fig. 3c), a trend for lower sucrose preference (Student’s *t*-test, *t*_14_=1.779, P=0.0970) (Fig. 3d), an increase in immobility time in the forced swim test (Student’s *t*-test, *t*_15_=2.537, P=0.0228) (Fig. 3D), and no difference in time spent in the open arms of the elevated plus maze (Extended Data Fig. 4a). In male mice, we did not observe any differences across behavioral tests (Fig. 3f-i, Extended Data Fig. 4b), supporting a female-specific increase in anxiety- and depression-like behaviors induced by neuronal expression of FEDORA in the mPFC, which mirrors the human female-specific increase in FEDORA levels in MDD.

**Fig. 3.**
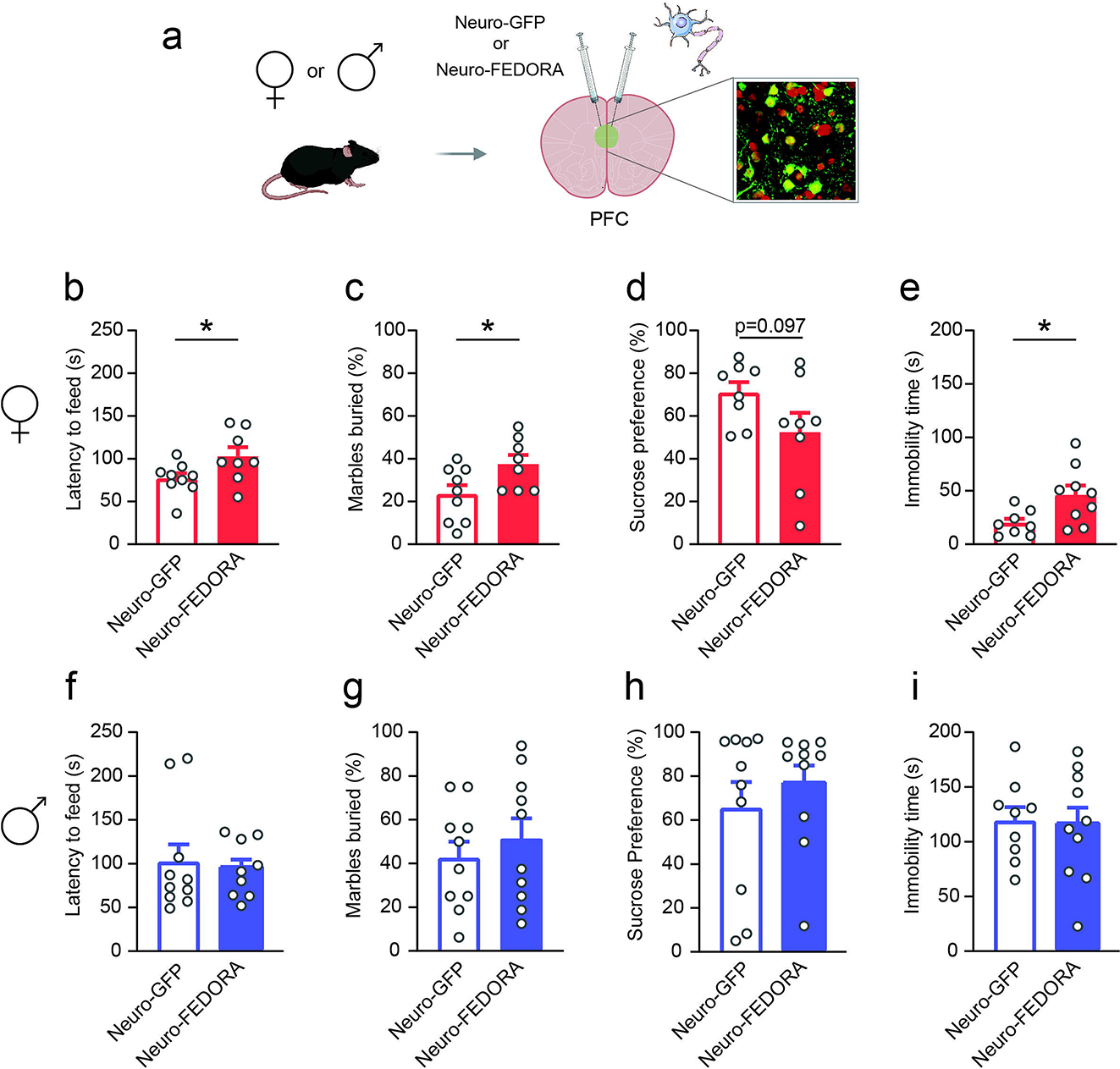
Expression of FEDORA in mouse mPFC neurons promotes anxiety- and depression-related behavioral abnormalities in females only. **(a)** Schematic representation of the experimental design. **(b-e)** In females, Neuro-FEDORA promotes increased latency to feed in the novelty suppressed feeding test, **(b)** increased number of marbles buried, **(c)** trend for deceased sucrose preference, **(d)** and longer immobility time in the forced swim test **(e)**. **(f-i)** No difference in any test was observed in male mice. N=8-10 group, * p<0.05. Bars represent mean ± SEM and dots represent individual data points.

Next, we expressed FEDORA in mPFC oligodendrocytes using a modified Adeno-associated virus (AAV) with a truncated version of human myelin-associated glycoprotein (*MAG*) promoter to guide cell-type specificity. We generated an AAV expressing FEDORA along with a GFP reporter (Oligo-FEDORA), or GFP only as control (Oligo-GFP) (Fig. 4a), and validated that this tool promotes FEDORA expression (Extended Data Fig. 3d), and confirmed that such expression predominates in oligodendrocytes (Extended Data Fig. 3f). We probed the sex-specific effect of Oligo-FEDORA in the mPFC of mice on anxiety- and depression-like behavioral phenotypes. We found that females treated with Oligo-FEDORA, compared to Oligo-GFP, showed increased immobility time in the forced swim test (Student’s *t*-test, *t*_15_=2.248, P=0.0401) (Fig. 4e), with no effect seen in the other tests in females (Fig. 4b-d, Extended Data Fig. 4c). Oligo-FEDORA had no effect in males across all tests used (Fig. 4 f-i, Extended Data Fig. 4d).

**Fig. 4.**
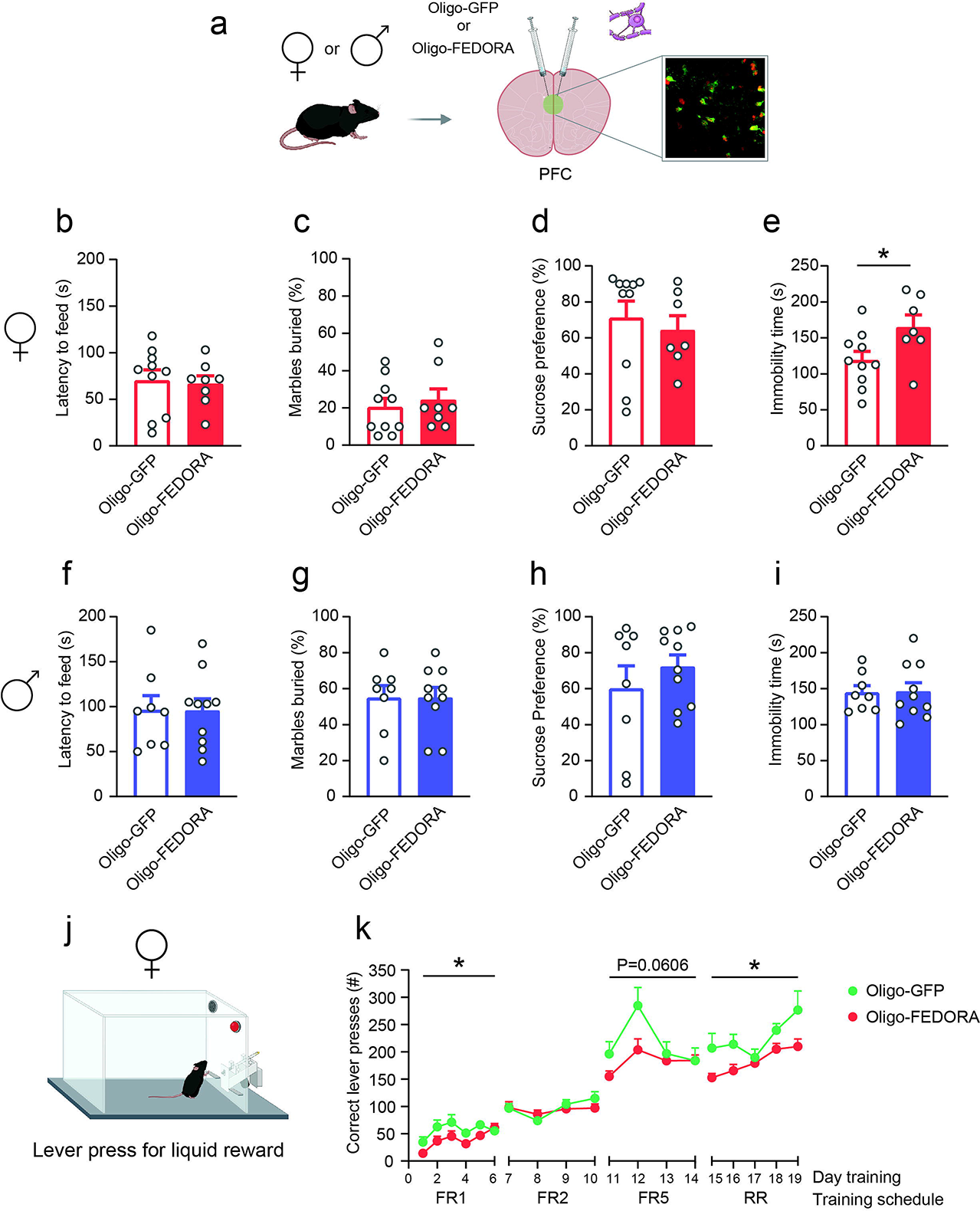
Expression of FEDORA in mouse mPFC oligodendrocytes promotes passive coping and attenuates reward learning in females only. **(a)** Schematic representation of the experimental design. **(b-e)** Bar graphs depicting that in females Oligo-FEDORA compared to Oligo-GFP does not alter behavior in the novelty suppressed feeding **(b)**, marble burying **(c)**, and sucrose preference **(d)** tests, but leads to increased immobility time in the forced swim test **(e)**. **(f-i)** No difference in any test was observed in male mice. N=8-10 group, * p<0.05. Bars represent mean ± SEM and dots represent individual data points. **(j)** Schematic representation of operant box used for training to lever press for liquid saccharin reward. **(k)** Graph representing that Oligo-FEDORA mice exhibited fewer correct lever presses than the Oligo-GFP group in the FR1 (Fixed Ratio) schedule and Random Ratio (RR) schedule, but not in the FR2 and FR5 schedule. N=10-12 per group, * p<0.05. Circles represent mean ± SEM.

To further explore the role of FEDORA in female mice in regulating motivation- and reward-related behaviors, we infused Oligo-FEDORA or Oligo-GFP into the mPFC of an additional cohort of mice, and trained them on an operant reinforcement task to press a lever to receive a liquid saccharin reward (Fig. 4j). Oligo-FEDORA treatment did not affect sucrose preference (Fig. 4d), suggesting that this manipulation does not affect the hedonic value of this reward. However, Oligo-FEDORA female mice demonstrated a lower number of correct lever presses compared to Oligo-GFP mice during the acquisition of responding for the saccharin reward on a fixed-ratio 1 (FR1; 1 response = 1 reward) schedule of reinforcement (Repeated measures two-way ANOVA, F virus_(1,20)_=4.837, P=0.0398) (Fig. 4k). Oligo-FEDORA mice reached similar levels of responding by the end of FR1 acquisition, and increasing the motivational demands of the task by increasing the schedule of reinforcement to FR2 or FR5 did not reveal a significant difference between the groups (Fig. 4k). However, switching to a more ambiguous random ratio (RR) 5 schedule of reinforcement, wherein each response has a 20% probability of saccharin delivery (5 responses per reward, on average), again revealed a deficit in performance in the Oligo-FEDORA mice (Repeated measures two-way ANOVA, F virus_(1,20)_=6.080, P=0.0228) (Fig. 4k). Together, these results indicate that FEDORA expression in mPFC oligodendrocytes impairs reward learning and performance in females.

### Effects of FEDORA Expression on Neuronal and Oligodendrocyte Function

To explore how neuronal FEDORA mediates its female-specific pro-depressive effects, we performed slice electrophysiological recordings from mPFC pyramidal neurons of mice of both sexes infected with either Neuro-FEDORA or Neuro-GFP as control. We found that neuronal FEDORA expression increased spontaneous excitatory postsynaptic current (sEPSC) amplitude (Fig. 5a,b) (Student’s *t*-test, *t*_13_=2.281, P=0.0401) and frequency (Student’s *t*-test, *t*_13_=2.137, P=0.0522) (Fig. 5a,c) in female but not male mice (Fig. 5f-h). Moreover, we found that Neuro-FEDORA compared to Neuro-GFP decreased the excitability of mPFC pyramidal neurons upon current injection in both female (Two-way ANOVA, F virus_(1,117)_=35.82, P<0.0001) (Fig. 5d,e) and male (Two-way ANOVA, F virus_(1,108)_=42.25, P<0.0001) (Fig. 5i,j) mice. Together, these results demonstrate both sex-specific and global effects of FEDORA expression on pyramidal neuron physiology in mPFC.

**Fig. 5.**
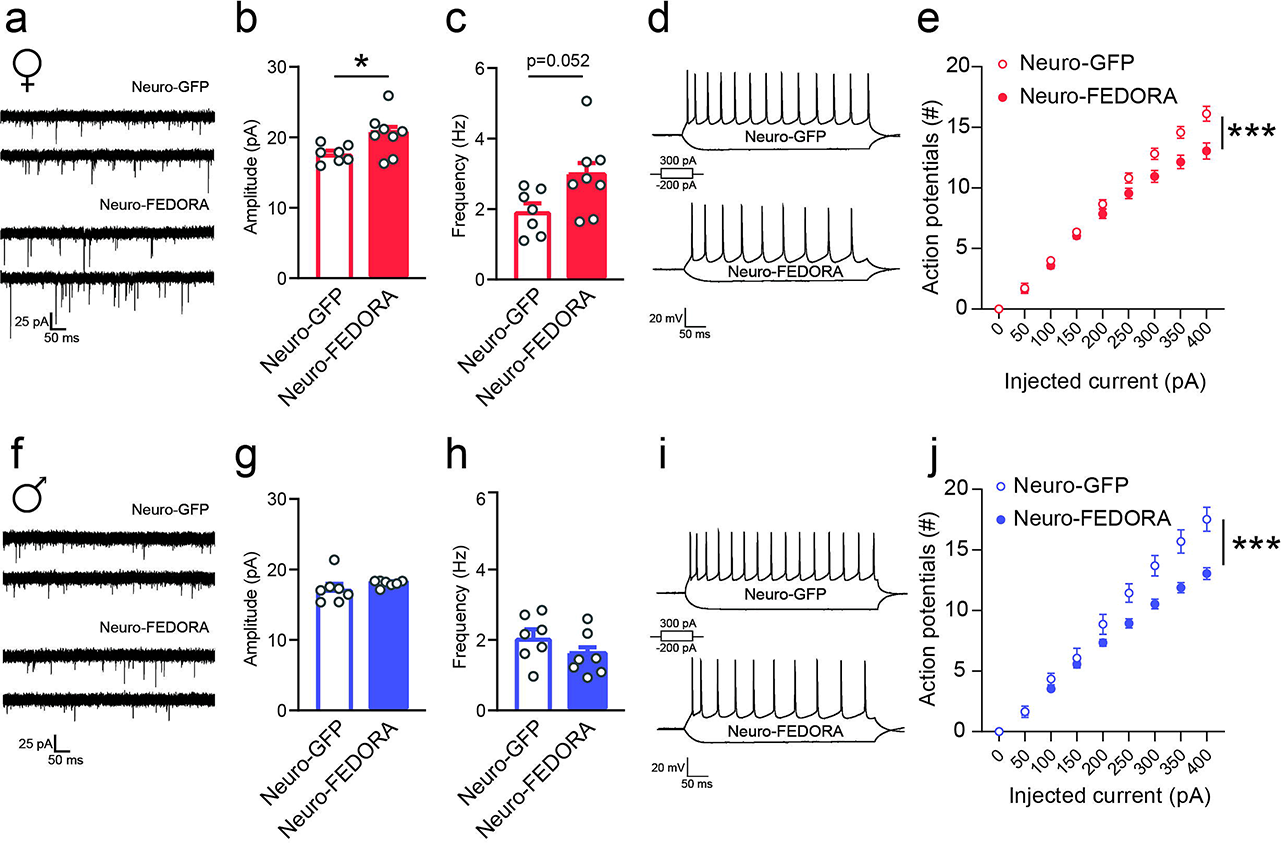
Neuronal FEDORA expression alters electrophysiological properties of mouse mPFC pyramidal neurons. **(a,f)** Representative tracks of spontaneous activity from female **(a)** and male **(f)** mice expressing Neuro-FEDORA or Neuro-GFP. FEDORA expression in female neurons increased sEPSC amplitude **(b)** and frequency **(c)**, with no change in males **(g,h)**. N=45-58 neurons from 7-8 mice per group, * p<0.05. Bars represent mean ± SEM and dots represent individual data points. FEDORA expression also reduces the excitability in female **(d,e)** and male **(I,j)** pyramidal neurons. N=45-58 neuron from 7-8 mice per group, *** p<0.001. Circles represent mean ± SEM.

Since MDD decreases myelin sheath thickness in PFC ^24, 25^, we hypothesized that this effect might be mediated by FEDORA function within oligodendrocytes. To test our hypothesis, we expressed Oligo-FEDORA or Oligo-GFP in mPFC of mice of both sexes and performed electron microscopy analysis of myelin thickness. We found that FEDORA expression decreased myelin thickness in mice of both sexes (Two-way ANOVA, F virus_(1,16)_=23.49, P=0.0002, post hoc Fisher LSD GFP:female vs FEDORA:female t_(16)_=3.034, P=0.0079; GFP:male vs FEDORA:male t_(16)_=3.820, P=0.0015) (Fig. 6) without changing axon diameter. This finding supports our hypothesis that FEDORA regulates myelination processes that are impaired in MDD. However, FEDORA expression in oligodendrocytes had no effect on the excitability of mPFC pyramidal neurons (data not shown), which together with findings with neuronal expression of FEDORA demonstrates cell-autonomous effects of FEDORA on both cell types.

**Fig. 6.**
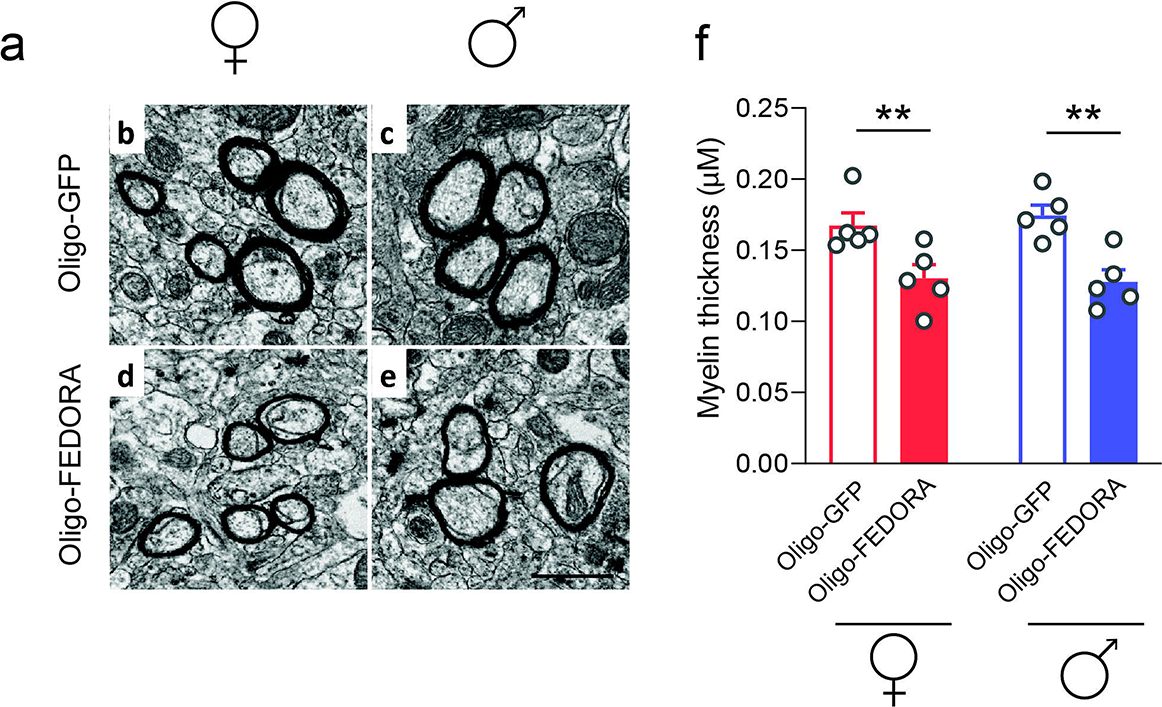
Oligodendrocyte FEDORA expression promotes myelin thinning in mouse mPFC. **(a)** Representative electron microscopy (EM) images from mouse mPFC infected with Oligo-FEDORA **(d,e)** or Oligo-GFP **(b,c)** of female **(b,d)** or male **(c,e)** mice. **(f)** Bar graphs represent quantification of the EM pictures and indicate thinner myelin in both male and female mice expressing FEDORA compared to GFP. N=91-126 axons from 5 mice per group, **p<0.01. Bars represent mean ± SEM and dots represent individual data points.

### Transcriptomic Effects of FEDORA Expression in Neurons and Oligodendrocytes

The ability of lncRNAs to act as epigenetic regulators of gene expression led us to the hypothesis that the sex- and cell-type-specific behavioral and cellular abnormalities induced by FEDORA are mediated by altered transcriptional regulation in the mPFC. Therefore, we characterized the female mouse mPFC transcriptome using bulk RNA-seq following the selective expression of FEDORA either in neurons or in oligodendrocytes. We found that the expression of FEDORA compared to GFP had a robust effect on gene expression, leading to 2837 differentially-expressed genes (DEGs; Fold Change>30% and q<0.05) when expressed in neurons (Fig. 7a) and 4037 DEGs when expressed in oligodendrocytes (Fig. 7f). As expected, cell-type analysis of the DEGs revealed significant enrichment for neuronal genes for DEGs induced with Neuro-FEDORA (Fisher Exact test P=2.18X10^-9^) and for oligodendrocyte genes induced with Oligo-FEDORA (Fisher Exact test P=7.72X10^-19^) (Fig. 7e). Notably, Oligo-FEDORA also significantly affected neuronal genes. Gene ontology analysis of the DEGs altered by Neuro-FEDORA highlights G-protein signaling and synaptic transmission (Fig. 7b), with translation processes implicated for Oligo-FEDORA (Fig. 7g).

**Fig. 7.**
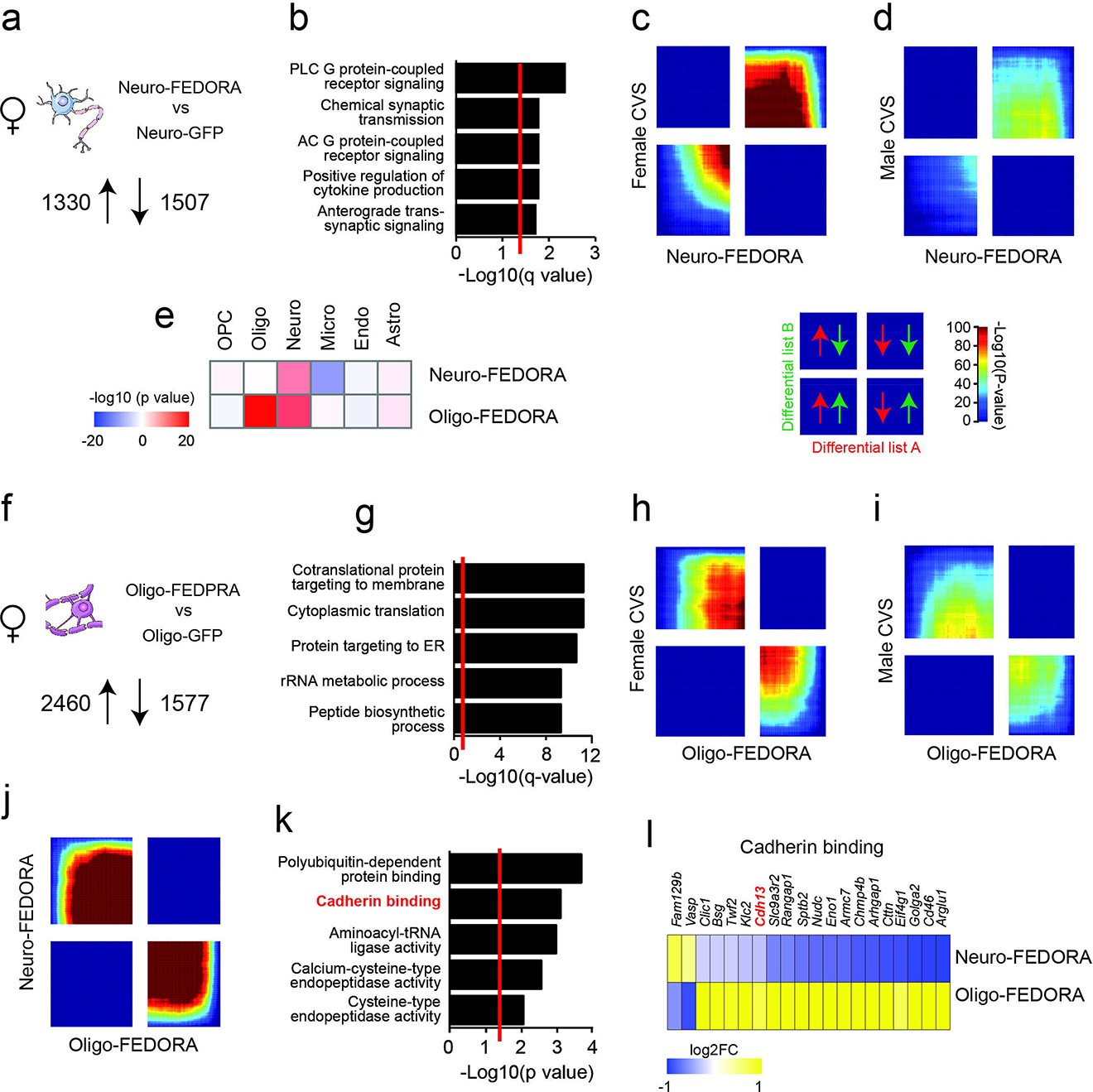
Opposing effects of FEDORA expression in neurons vs oligodendrocytes on gene expression. **(a,f)** Schematic representation of bulk RNA-seq experiments in mPFC of female mice upon the selective expression of FEDORA or GFP in neurons **(a)** or in oligodendrocytes **(f)**. The numbers indicate the number of genes either up-or downregulated by FEDORA expression compared to GFP controls. N=4-5 samples per group. **(b, g)** Bar graphs indicating the top biological process GO ontology terms enriched in genes differentially expressed by Neuro-**(b)** or Oligo-FEDORA **(g).** The red lines indicate the significance threshold at q<0.05. **(c,d,h,i)** Heatmaps from a rank rank hypergeometric overlap (RRHO) analysis ^27^ comparing the effects of FEDORA expression in mPFC to that of chronic variable stress (CVS) on the mPFC transcriptome of mice of both sexes. Neuro-FEDORA transcriptional signature overlaps more strongly with females **(c)** than males **(d)** exposed to CVS for both co-down and co-upregulated signals. Oligo-FEDORA displays an opposing pattern of overlap with the transcriptional signature of CVS in females **(h)** more so than males **(i)**. **(e)** Heatmaps showing significant positive enrichment for neuronal markers in the genes differentially expressed by Neuro-FEDORA, and for oligodendrocytes and to lesser extent neurons by Oligo-FEDORA. Astro = Astrocytes; Endo = Endothall; Micro = Microglia; Neuro = Neurons, Oligo = Oligodendrocytes; OPC = Oligodendrocyte progenitor cells. **(j)** Heatmaps from RRHO analysis showing opposite effects on the transcriptome by Oligo-FEDORA compared to Neuro-FEDORA. **(k)** Bar graphs indicating the top biological process GO ontology terms enrichment analysis for genes that have opposing regulation by Oligo-FEDORA compared to Neuro-FEDORA. (**l**) Heatmaps highlighting the opposing regulation of the Cadherin binding genes opposingly regulated by FEDORA in the two cell types, including CDH13 the antisense gene of FEDORA.

Next, we compared the effects of FEDORA on the mPFC transcriptome to those of chronic variable stress (CVS), which induces a similar range of anxiety- and depression-like behavioral abnormalities in female and male mice (data from ^26^). We utilized rank-rank hypergeometric overlap (RRHO) analysis which compares differential expression across two datasets in a threshold-free manner ^27^. We found that the Neuro-FEDORA vs. Neuro-GFP transcriptional signature exhibits a robust overlap with downregulated genes of female mice exposed to CVS compared to non-stressed controls (Fig. 7c), but this was not observed for males (Fig. 7d). Interestingly, when we performed the same analysis on the Oligo-FEDORA vs. Oligo-GFP signature we found an opposite effect that was again more pronounced in females vs males: a significant subset of genes that were upregulated by FEDORA expression in oligodendrocytes were downregulated by CVS and, conversely, genes that FEDORA downregulated are upregulated by CVS (Fig. 7h-i).

We further probed this opposite cell-type-specific transcriptional effect of FEDORA and found significant enrichment for genes with a baselines sex difference in mouse mPFC (from our previous publication ^26^) within the genes regulated by Neuro-FEDORA (Fisher Exact test P=2.303X10^-14^) and Oligo-FEDORA (Fisher Exact test P=0.202) (Extended data Fig. 5a). Specifically, numerous genes that are expressed at higher levels at baseline in females vs males are downregulated upon Neuro-FEDORA expression (Extended data Fig. 5b). Moreover, we found a cell-type-specific effect of FEDORA on the expression pattern of the 360 PCGs that significantly correlate with FEDORA in our female human postmortem brain dataset (Extended data Fig. 5c). Next, we directly compared the Neuro-FEDORA to the Oligo-FEDORA transcriptional profiles and found a strong inverse effect on gene expression between the cell types (Fig. 7j). Gene ontology analysis of the genes demonstrating this opposite regulation highlights protein binding, particularly cadherin binding (Fig. 7k). One of these genes is *Cdh13* (Fig. 7l), the gene to which FEDORA is antisense. Together, these results suggest that FEDORA promotes opposing cell-type-specific patterns on gene expression and implicate its host gene, *Cdh13*, as one potential mediator of these effects.

### FEDORA Levels in Human Blood Track MDD Symptoms in Females

Finally, we tested the translational potential of the sex-specific regulation of FEDORA by testing whether its circulating levels may serve as a diagnostic or prognostic biomarker for female MDD. We probed total blood levels of FEDORA in a cohort of MDD cases of both sexes compared to matched HCs. The cases were treatment-resistant patients enrolled in a study in which they were treated with the rapidly-acting antidepressant ketamine, and blood was re-collected 24 h after treatment (Fig. 8a). We performed qPCR to probe the levels of FEDORA and found that its expression levels were higher in females with MDD compared to HCs before treatment (Two-way ANOVA, F group_(2,100)_=4.523, P=0.0132, post hoc Fisher LSD HC:female vs MDD pre:female t_(100)_=2.666, P=0.0089) (Fig. 8b). This effect was not apparent in males. Ketamine treatment did not normalize the levels of FEDORA at the group level. However, we found a significant correlation between the change in FEDORA expression at pre-to post-treatment time points and the change in depression severity as measured by the Quick Inventory of Depressive Symptomatology (QIDS) in female MDD subjects (Pearson correlation, r=0.605; P=0.0377) (Fig. 8c), but not in males (Fig. 8d). Together, these results suggest that FEDORA total blood levels may serve as a female-specific diagnostic tool for MDD which tracks treatment response to ketamine in females.

**Fig. 8.**
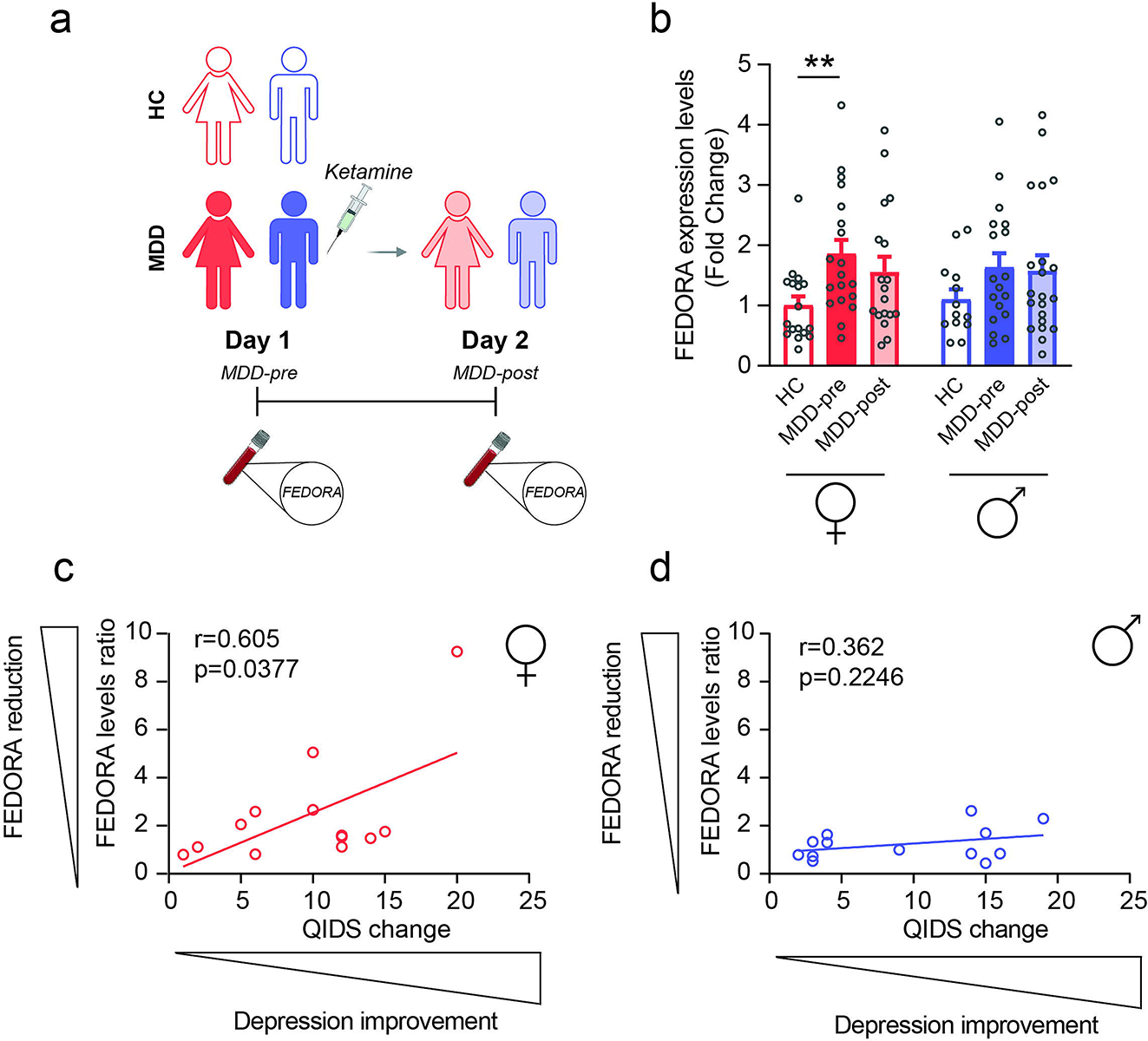
Blood FEDORA levels are associated with MDD symptoms and response to ketamine treatment in females only. **(a)** Schematic representation of the experimental design. Blood was collected from treatment-resistant MDD (MDD-pre) and HC subjects of both sexes. MDD subjects were treated with the fast-acting antidepressant ketamine and blood was drawn again one day later (MDD-post) along with clinical assessment. **(b)** Bar graph showing that blood levels of FEDORA as measured by qPCR are higher in MDD-pre females compared to HCs, but not in males. N=13-21 per group, **p<0.01. Bars represent mean ± SEM and dots represent individual data points. **(c-d)** Graphs demonstrating the correlation between the change in MDD-post vs MDD-pre in FEDORA levels and the change in the Quick Inventory of Depressive Symptomatology (QIDS) scale. There is a significant positive correlation between the improvement in MDD and the reduction of FEDORA levels in female **(c)** but not male **(d)** subjects. N=15-16 per group.

## Discussion

Here we report that lncRNAs exhibit a striking sex difference in expression across the brain of healthy individuals, and that this sex-specific pattern of expression is broadly lost in patients diagnosed with MDD. We highlight one such lncRNA, which we named FEDORA, after demonstrating that it is upregulated across cortical brain regions in females, but not males, with MDD compared to HCs in two independent cohorts. By expressing FEDORA selectively in neurons or in oligodendrocytes of mouse mPFC, we validated its role as a sex-specific mediator of anxiety- and depression-like behavior. We also demonstrate a cell-type-specific influence of FEDORA on neuronal and oligodendrocyte function: FEDORA alters synaptic physiology when expressed in neurons with no effect seen upon expression in oligodendrocytes; FEDORA alters myelin thickness in oligodendrocytes; and FEDORA promotes opposite transcriptional programs in these two cell types. Interestingly, the transcriptional effects of expressing FEDORA in neurons and oligodendrocytes implicate regulation of the cell adhesion molecule CDH13 to which FEDORA is an antisense transcript. Finally, we show that blood levels of FEDORA reflect depression diagnosis specifically in women and correlate with the extent of their response to ketamine treatment. Taken together, these results highlight the role of lncRNAs in maintaining the transcriptional homeostasis underlying sex differences in the brain and point to FEDORA as a diagnostic and therapeutic target for depression in women.

Sex differences have emerged as critical mediators of stress, anxiety, and depression in recent years ^28^, and several lines of evidence suggest that males and females exhibit mostly distinct patterns of transcriptional regulation throughout limbic brain regions in MDD in humans and after chronic stress in mice ^26, 29,30,31^. Here, we report that MDD erases transcriptomic sex differences typically observed in HCs across brain regions associated with depression. This phenomenon is similar to what we observed in the brains of adult mice that were exposed to social isolation in adolescence ^32^. We note that the loss of sex differences in MDD is more pronounced for lncRNAs compared to PCGs. There are reports of a sex-specific expression pattern of lncRNAs in mammalian gonads ^22^ and mouse liver ^33, 34^, and in the brain of zebrafish ^35^ and drosophila ^36^. Specific lncRNAs such as the maternally imprinted *Peg3* were linked to regulating behaviors and gene expression ^37^, whereas *Xist* is associated with female-specific X chromosome inactivation^38^ and sex-specific gene expression silencing ^39^. There are reports that sex-biased lncRNA expression in the liver associates with sex-biased accessible chromatin regions and hormonal binding sites ^33^, however, the processes that guide baseline sex differences in lncRNA expression in the brain remain unknown, particularly as relates to altered expression patterns in MDD.

FEDORA is antisense to *Cdh13* mRNA which encodes a member of the cadherin superfamily of cell adhesion molecules and is expressed predominantly in neurons as well as in oligodendrocyte progenitor cells and mature oligodendrocytes ^40^. In neurons, CDH13 is localized to the cell membrane, where it acts as a negative regulator of axon growth during neuronal differentiation. Genetic studies have linked variations in *CDH13* to diagnosis with attention deficit hyperactivity disorder, alcohol dependence, and MDD ^41,42,43,44^. Moreover, a recent proteome-wide association postmortem brain study of MDD reported CDH13 as a significant hit ^45^. Notably, another cadherin family member, PCFG19, has been suggested to exhibit a female-specific role in synaptic and cognitive function ^46^. While CDH13 is not regulated in MDD at the RNA level in our postmortem human brain dataset, nor in mouse models of chronic stress ^26, 47^, we found a robust opposite regulation of *Cdh13* upon expression of FEDORA in neurons vs oligodendrocytes. These findings now warrant further investigation of CDH13 as a potential mediator of some of FEDORA’s effects on the brain under normal and pathological conditions.

We report here that neuronal expression of FEDORA in mPFC promotes anxiety- and depression-like behavioral abnormalities. Our findings align with a large body of literature indicating that neuronal dysfunction in the PFC, the central hub of executive functions, contributes importantly to MDD. There are reports from postmortem brain studies from MDD subjects ^48, 49^ as well as rodent chronic stress models ^47, 50, 51^ describing the loss of synapses, a decrease in synaptic-related genes, and alterations in mPFC neuronal electrophysiological properties. Similarly, we found that neuronal expression of FEDORA promotes altered expression of synaptic genes along with changes in neuronal excitability. Notably, our current results that FEDORA-induced behavioral abnormalities in females coincides with increased amplitude and frequency of sEPSCs of mPFC neurons are in line with our former study of another lncRNA, LINC00473, which promotes females-specific stress resilience and opposite effects on sEPSCs ^17^, and with reports on sex- and pathway-specific effects of chronic stress on mPFC physiology ^52^. Collectively, our study adds FEDORA as a mediator of the female brain’s molecular arsenal that controls mPFC excitability and that is aberrantly expressed in MDD.

Multiple studies have found deleterious effects of MDD on oligodendrocytes and myelination in the PFC (reviewed in ^24, 53, 54^), reporting lower density of oligodendrocytes in MDD cases ^55, 56^, white matter abnormalities, and downregulation of oligodendrocyte genes ^57,58,59,60^. Rodent studies using several chronic stress models report reduced oligodendrocyte proliferation and thinning of myelin in the PFC, which was reversed by antidepressant treatment ^61,62,63,64,65,66,67^. Consistent with these findings, we show that FEDORA expression in oligodendrocytes produced a pro-depressive-like behavioral effect, although the behavioral abnormalities induced by oligodendrocyte FEDORA expression are different from those induced by neuronal FEDORA expression. FEDORA expression in oligodendrocytes also caused thinning of myelin in the mPFC. Notably, while the pro-depressive behavioral effects were only apparent in female mice, we noted myelin thinning in both sexes. This lack of sex specificity aligns with the decreased intrinsic (as opposed to synaptic) excitability we report in neurons from mice of both sexes upon FEDORA expression in neurons. We speculate that these two phenomena may be interrelated, as decreased myelin thickness can alter neuronal excitability ^68,69,70^. As well, we found enrichment for altered expression of synaptic and postsynaptic genes when expressing FEDORA in female mice in neurons (Extend Fig. 5d) or in oligodendrocytes (Extend Fig. 5e). We speculate that reciprocal interactions between oligodendrocytes and neurons support dynamic plasticity in the thickness of myelin sheaths regulated by FEDORA, potentially through its interaction with CDH13. Alternatively, in line with evidence for aberrant chromatin structure in oligodendrocyte progenitor cells within the PFC of chronically stressed mice ^66^, we hypothesize that FEDORA may regulate chromatin structure, as often is the case with lncRNAs, to direct transcriptional processes leading to pro-depressive effects.

We also report that the sex-specific differential regulation of FEDORA in the brains of MDD patients is observed in the circulation, with higher FEDORA levels observed in whole blood of treatment-resistant depressed females, but not males. Although previous studies have profiled the genome-wide transcriptional changes in blood of individuals with MDD compared to HCs ^71,72,73^, none have included sex as a biological variable. We also identified a correlation between reductions in FEDORA blood levels by treatment with ketamine and its therapeutic efficacy. Interestingly, while rodent studies report heightened sensitivity to ketamine in females (reviewed in ^74, 75^), meta-analyses of human studies did not find sex differences in the effects of acute ketamine treatment for MDD ^76, 77^. It is unclear if the mechanisms which lead to upregulation of FEDORA in the brain of depressed women are similar to those that alter its expression in the circulation. Interestingly, FEDORA is expressed predominantly in antibody-producing B cells in blood^78^. Most studies to date on circulating factors in the context of stress and depression focus on the innate immune system or T-cell-related processes and not on B-cells ^79^, therefore, our unique finding of a B-cell-specific depression biomarker is interesting and warrants further investigation.

To conclude, our findings support the view that human-specific lncRNAs serve a prominent regulatory role in higher brain function, including mood, under normal and pathological conditions. We acknowledge that mechanistic studies of the role of a human-specific transcript like FEDORA in the context of a complex brain disorder presents many challenges. However, even though mice do not express FEDORA or any apparent homologue, we hypothesized that FEDORA would nevertheless produce specific effects in mice based on the knowledge that many lncRNAs act by binding to DNA, RNA, or protein targets and that mice likely express many of FEDORA’s targets even if they do not express FEDORA itself. Indeed, the dramatic sex- and cell-type-specific effects induced by FEDORA expression in neurons or in oligodendrocytes support this hypothesis. Additionally, our approach of expressing a human-specific lncRNA in mice has successfully been used by us and other groups to understand the actions of other lncRNAs in brain and other tissues ^17, 80, 81^. We speculate that human-specific lncRNAs such as FEDORA arose later in evolution as an additional layer of regulation of increasingly complex molecular pathways already established in earlier mammals such as mice. Our findings with FEDORA support this possibility and elaborate novel molecular mechanisms that contribute to sex differences in higher brain function at baseline and in the context of MDD.

## Online Methods

### Animals

Experiments utilized eight-week-old C57BL/6J male or female mice (Jackson Laboratory). Mice were habituated to the animal facility for 1 week before any manipulations and were maintained at 23–25°C with a 12 h light/dark cycle (lights on 7:00 A.M.) with ad libitum access to food and water. Experiments were conducted following the guidelines of the Institutional Animal Care and Use Committee (IACUC) at Mount Sinai.

### Behavioral testing

#### Novelty suppressed feeding

The novelty suppressed feeding test assesses anxiety- and depression-related behaviors^82^. Mice were food-deprived for 24 h before testing and habituated to the experimental room for 1 h before testing. Mice were placed in the corner of a 50 × 50 × 20 cm box covered with bedding with a single food pellet placed in the center of the arena. The latency to eat was recorded under red light conditions during 5 min of testing. Mice were then transferred to their home cage and their latency to eat was recorded there as well.

### Sucrose preference

Mice were housed individually and given a choice between two 50-mL conical tubes with either water or 1% sucrose solution for 24 h ^83^. The bottles were weighed, and sucrose preference was calculated by determining the percentage of sucrose consumption divided by total liquid intake over one day. Total liquid intake did not differ across experimental groups studied.

### Elevated plus maze

For the elevated plus maze, which assesses anxiety-related behavior ^84^, mice were acclimated to the testing room for 1 h before testing. Animals were tested in a cross-shaped maze consisting of two 12 × 50 cm open arms and two closed arms with 40 cm high walls under red light for 5 min. Behavior was tracked using an automated system (Ethovision, Noldus).

### Marble burying

The marble burying test for anxiety-related behaviors ^85^ was conducted under red light conditions. After one h of acclimation to the testing room, mice were put in a standard mouse cage filled with 15 cm of corn cob bedding topped with 20 glass marbles. After 15 min, the mice were removed and the number of marbles fully or partiality buried were counted.

### Forced swim test

The forced swim test, which assesses an animal’s response to an acute stress, was performed after 1 h of habitation to the testing room. Mice were tested in a 4 L glass beaker, containing 3 L of water at 25±1°C for 6 min under white light conditions. The behavior was video tracked using an automated system (Ethovision, Noldus) ^83^.

### Operant lever press task

Mice were trained to lever-press for a liquid saccharin reward in operant conditioning boxes (22 × 18 × 13 cm; Med Associates, St Albans, Vermont) as described previously ^86^. Each box contained two levers and a motor-driven dipper that delivers 0.02 ml of 0.2% saccharin through a hole in the magazine floor per correct lever press. Operant boxes were illuminated and enclosed in a sound-attenuating chamber equipped with a ventilation fan. Training started one month after surgery that delivered Oligo-FEDORA or Oligo-GFP to the mPFC. Female mice were acclimated to water restriction and saccharin for three days in their home cage and then maintained on water restriction (2 h *ad libitum* access per day) throughout testing. First, mice were trained to retrieve saccharin from the reward magazine when it was presented. During the training, the shift to the next task or schedule was carried out after the behavior stabilized. Next, in 30-min daily sessions, mice were trained to lever press for saccharin on a fixed ratio 1 (FR1) schedule of reinforcement, in which each correct lever press is rewarded, for 6 days. This was followed by a fixed ratio 2 (FR2) for 4 days, a fixed ratio 5 (FR5) for 4 days, and finally a schedule of random ratio 5 (RR) reinforcement with a 1-in-5 chance of the response being rewarded for 5 days.

### Viral vectors

For neuronal expression FEDORA was synthesized (Gene Script) and Gateway (Invitrogen) cloned into a p1005+ HSV vector. This vector expresses eGFP with a CMV promoter while the gene of interest is driven by the IE4/5 promoter. Empty p1005+ expressing eGFP only was used as the control. The plasmids were packaged into HSVs that are highly neurotrophic ^87^ and display rapid induction, therefore, experiments were performed 3 days after infection. For oligodendrocyte expression, FEDORA was subcloned into pAM/0.3KB-MAG-eGFP AAV vector, contributed by Matthias Klugmann (University of New South Wales (Australia). The plasmid includes a 0.3 kB truncated form of the promoter of the human myelin-associated glycoprotein (*MAG*) gene that drives transgene expression in oligodendrocytes ^88^ and fits within AAV plasmid size constraints. Empty pAM/0.3KB MAG-eGFP expressing eGFP only was used as the control. The plasmids were used to make AAV1 serotype at the vector core of the University of Pennsylvania at 10^13^ GC/ml concentration. Experiments using this AAV tool were performed one month following mPFC infection when transgene expression in maximal. Validation of HSV and AAV vectors was confirmed by epifluorescence microscopy and qPCR.

### Stereotactic surgery

Mice were anesthetized with a mixture of ketamine (100 mg/kg) and xylazine (10 mg/kg), and positioned in a small-animal stereotactic instrument (Kopf Instruments). The skull surface was exposed, and 33-g syringe needles (Hamilton) were used to bilaterally infuse 0.5 µl of viral vectors at a rate of 0.1 µl/min. mPFC coordinates relative to the Bregma were: AP + 1.8 mm; ML +0.75 mm and DV −2.7 mm with 15° angle.

### Electrophysiology

Pyramidal neurons in mPFC of male and female mice were studied in brain slices one day after mice received intra-mPFC injections of Neuro-FEDORA or Neuro-GFP. Virally infected neurons were distinguished from non-infected neurons by their GFP signals under epifluorescence microscopy. Spontaneous excitatory postsynaptic potential EPSCs (sEPSCs) were recorded for 3 min in the voltage-clamp mode, with a K-based internal solution (in mM: 130 K-methanesulfonate, 10 KCl, 10 HEPES, 0.4 EGTA, 2.0 MgCl2, 3 MgATP, 0.5 Na3GTP, 7.5 phosphocreatine, pH 7.4; 285 mOsm). The frequency and amplitude of sEPSCs were analyzed with Minianalysis Program (Synaptosoft). Parallel experiments were performed on mPFC pyramidal neurons 30 days after intra-mPFC injection of Oligo-FEDORA or Oligo-GFP.

### Electron Microscopy

Male and female mice were perfused as described previously ^66^ for electron microscopy (EM) analysis one month after stereotactic delivery of Oligo-FEDORA or Oligo-GFP (N=5 per group). Brains were harvested and vibratome sectioned into 60 μm slices containing the mPFC. Sections were resin embedded, thin sectioned at 90 nm, stained with uranyl acetate and lead citrate, and mounted on 200 mesh copper grids. Ten images at 10,000X were collected per mouse using a transmission electron microscope JEOL JEM 1400Plus equipped with a Gatan CCD camera. ImageJ was used to measure both axon caliber and myelin fiber diameter for about 100 myelinated axons per mouse. All analyses were performed blind to the experimental conditions.

### Immunohistochemistry, Imaging, and Cell Counting

Adult mice were terminally anesthetized with ketamine/xylazine, transcardially perfused with PBS followed by 4% paraformaldehyde, and brains were postfixed in 4% paraformaldehyde at 4°C. Coronal sections (30 µm) were sliced on a vibratome, washed 3x with PBS 0.1%Triton-X (PBST), and blocked with normal donkey serum (3%) at room temperature for 1 h in PBST. Sections were then incubated in primary antibodies overnight at 4°C in blocking solution (1:2000 Chicken Anti-GFP, # GFP-1020, Aves Lab; 1:500 Rabbit Anti-Olig 2, #AB9610, Abcam; Rabbit Anti-GFAP #ab7260, Abcam; Rabbit Anti-Neun # NBP1-77686, Novus). Sections were then washed 3x in PBS and incubated with secondary antibodies (1:500 Alexa 488 Donkey Anti-Chicken # 703-546-155, Jackson ImmunoReserach; 1:500 Cy3 donkey anti-rabbit #711-166-152, Jackson ImmunoReserach) for 90 min at room temperature. Sections were washed in PBS, mounted on slides and coverslipped with ProLong Gold with Anti-fade including Dapi (Invitrogen). For quantification of immunohistochemistry, slides were imaged on a Zeiss LSM710 confocal microscope. Only high-quality slices at a similar level along the A-P axis were imaged at 20x-magnification, such that 3-4 slices per mouse were imaged and quantitated (n=3-5 mice per marker). Analysis of cell counts and colocalization was performed using ImageJ by an observer blind to condition. A mean count per mouse was used for statistical analysis.

### Human postmortem situ hybridization

Fluorescent in situ hybridization (FISH) was performed on an independent cohort of male and female MDD cases (N=10 per group) (Extended Data Table 1). The samples were obtained by The University of Texas Southwestern Medical Center at Dallas with consent from the next of kin. Collection criteria included <24-h postmortem interval (PMI), and no direct head trauma or other medical illness. The brains were taken from the skull, put on ice, dissected, and flash frozen. Samples were stored at −80°C in plastic under a vacuum seal. The diagnosis was performed based on hospital and medical records and interviewer with the next of kin. This information was reviewed by four clinicians that came to a consensus diagnosis using DSM-IV criteria. Subjects were matched to controls for age, pH, RNA integrity number (RIN), and PMI.

FISH was performed using RNAscope in human rostral anterior cingulate cortex (rACC) in layers II-VI for gray matter and from subcortical white matter. Fixed-frozen tissue samples were cryostat sectioned (14 µm) and mounted on SuperFrost Plus slides. RNAscope Multiplex Fluorescent reagent kit and probes for FEDORA (LOC105371366) as well as RBFOX3 (neuronal marker), OLIG2 (oligodendrocyte makers), and fluorophore (TSA) were used according to manufacturer instructions. Briefly, the slides were subjected to pretreatment, including hydrogen peroxide, epitope retrieval in 100°C water bath, and digestion enzyme. Next, the sections received FEDORA probes and were hybridized for 2 h at 40°C. Amplification and detection were performed using TSA Fluorescein (1:750) before counterstain with Dapi. Confocal images were acquired using a Zeiss LSM800 confocal microscope. High-resolution and high-magnification tiling images were taken from the region of interest (ROI). Quantification was performed with Fiji/ImageJ (NIH). The number of puncta was extracted, and density was calculated by dividing the size of the ROI.

### RNA extraction

To collect FEDORA expressing mPFC samples, mice were cervically dislocated and the brains were rapidly removed. Bilateral 1 mm-thick 14 g microdissections of mPFC were taken from fresh slices and flash-frozen on dry ice. Total RNA was isolated with QIAzol (Qiagen) and purified with the RNeasy Micro Kit (Qiagen), including on-column DNase treatment. For human subjects’ blood, intravenous blood was collected using EDTA tubes and stored in a −80°C. Total RNA was extracted using the total RNA Purification Kit (Norgen Biotek) including on-column DNase treatment. All samples were tested for concentration and purity using a NanoDrop (Thermo Fisher). Samples used for RNA-seq were analyzed using a bioanalyzer RNA Nano chips (Agilent) for integrity.

### RT and qPCR

RNA amount was normalized across samples and cDNA was created using iScript (Bio-Rad). Real-time PCR reactions were run in triplicate and SYBR-green was used in a QuantStudio 7 (ThermoFisher) qPCR machine. The 2^-ΔΔCt^ method was used to calculate relative gene expression normalized to controls with *Hprt1* as the house-keeping gene. See Extended Data Table 3 for primer sequences.

### Sequencing library preparation

RNA libraries were prepared from female mouse mPFC infected with Neuro-FEDORA or Neuro-GFP, as well as from Oligo-FEDORA or Oligo-GFP. The successful manipulation of FEDORA was confirmed in the samples used for sequencing by qPCR. Libraries were made using 1 µg of purified RNA for neuron sequencing using the ScriptSeq Complete Kit (Epicentre) and 500 ng RNA for oligodendrocyte with the TruSeq Stranded Total RNA (Illumina). Library preparation kits were changed due to discontinuous production of the Epicentre kit. For library prep, the cDNA was synthesized from ribosomal depleted RNA and then fragmented total RNA using random hexamers, followed by terminal tagging. The libraries were PCR amplified and then purified using AMPure XP beads (Beckman Coulter). Barcode bases were introduced at the end of the adaptors during PCR amplification steps. Quality and concentration of libraries were confirmed on a Bioanalyzer (Agilent), and libraries were sequenced on an Illumina Hi-seq machine with 150-bp paired-end reads by Genewiz. Samples were multiplexed to produce >50 million reads per sample. Sample size was N=4-5 independent samples per group.

### RNA-seq analysis of human postmortem brain and mouse brain

Human postmortem RNA-seq data from MDD and HC were collected, analyzed, and reported as part of a published study ^26^. Briefly, brain tissue was obtained from the Douglas Bell Canada Brain Bank Québec. Males and females were group-matched for age, pH, and PMI. Inclusion criteria for both cases and controls were French-Canadian European origin and sudden death. Forty-eight subjects were recruited, and tissue was collected from six brain regions: the orbitofrontal cortex (OFC; BA11), dorsolateral PFC (BA8/9; dlPFC), cingulate gyrus 25 (BA25; cg25; vmPFC), anterior insula (aINS), nucleus accumbens (NAc), and ventral subiculum (vSUB). The study was approved by the research ethics boards of McGill University.

Sex differences were studied based on the differential expression analyses between HC females and males by brain region which was part of the original RNA-seq study ^26^. Here, we filtered these lists to identify genes with baseline sex differences with fold change >30% and p<0.05. We define a loss of baseline sex differences if an RNA that is upregulated in HC females compared to males is downregulated in females with MDD compared to female HC (fold change >30%) or upregulated in male MDD compared to male HC. Or the inverse patterns, RNA which is downregulated in HC females compared to males is upregulated in females with MDD compared to female HC or downregulated in male MDD compared to male HC. We also only included lncRNAs in our analyses that lost their baseline sex difference only in one of the sexes and not in both.

The correlation analysis between lncRNA and PCG expression data was part of our former study ^17^, where we performed genome-wide Pearson correlation between the brain site normalized expression level (logCPM) of a lncRNA and PCG for each sex separately. Permutation analysis of the same number of random genes iterated 10,000 times was used to calculate a threshold for significant Pearson correlation at R>0.58771 or R<-0.59516, with 23% FDR.

For RNA-seq differential expression analysis, raw reads were aligned to the mouse genome (mm10) using HISAT2 ^84^. The average mapping rates for neurons and oligodendrocytes were 84% and 89%, respectively. Quality control was performed using FastQC ^5^ and principal component analysis was used to identify outliers. Normalization and differential expression analysis were performed using DESeq2 ^89^. Raw data were deposited in GEO (GSE188930). Significance was set at FDR correct q<0.05 and a fold-change (FC) threshold was set at >1.3. See Extended Data Table 4 for Neuro-FEDORA and Extended Data Table 5 for Oligo-FEDORA differential gene expression lists. Heatmaps were generated using Morpheus (https://software.broadinstitute.org/morpheus). Gene ontology analysis for enrichment of biological process was performed using Enricher ^90^ and SynGo was used to analyze synaptic genes enrichment ^91^. Cell type enrichment analysis was performed by comparing the differentially expressed gene (DEG) signature to a mouse brain cell type-specific marker lists ^92^ followed by a Fisher Exact test. Rank-rank hypergeometric overlap (RRHO) analysis was used to evaluate the overlap of differential expression lists between Neuron-or Oligo-FEDORA expression and mouse chronic variable stress (CVS) signature from our published dataset ^26^. A two-sided version of this analysis was used to test for coincident and opposite enrichment ^27^.

### Human blood study

Male and female participants between the ages of 18–65 were recruited through the Depression and Anxiety Center for Discovery and Treatment at the Icahn School of Medicine at Mount Sinai for a ketamine study ^93^. The diagnosis was assessed using the structural clinical interview for the Diagnostic and Statistical Manual of Mental Disorders-Fifth Edition (DSM-V). Participants met diagnostic criteria for a current major depressive episode of at least moderate severity as measured by the Clinical Global Impression-Severity scale (CGI-S), as well as a lifetime history of non-response to at least two trials of antidepressants. Demographic information (Extended data Table 2), as well as an additional psychiatric evaluation, was collected within 28 days before the initial blood draw. Exclusion criteria included a lifetime history of a psychotic disorder, bipolar disorder, alcohol or other substance abuse (confirmed by urine toxicology), any unstable medical condition, any systemic inflammatory or autoimmune disease, current antidepressants use, pregnancy or nursing. MDD severity at baseline was measured using the Quick Inventory of Depressive Symptomatology (QIDS) after which blood samples were collected. Thereafter, participants received a single ketamine I.V. infusion of 0.5 mg/kg over 40 min. One day after the infusion, both MDD severity scores and blood samples were collected again. The study was approved by the program for the protection of human subjects at Mount Sinai and participants provided informed consent.

### Statistical analyses

All statistics were performed using Prism 9. Outliers were defined as more than 2.5 standard deviations from the group mean and were removed from the analysis. Main effects and interactions were determined using one-or two-way ANOVA with Fisher LSD post hoc analysis. Student’s t-test was used for comparisons of only two groups. All significance thresholds were set at p<0.05.

## Acknowledgements

This work was funded by grants from the National Institute of Mental Health (R01MH051399) and the Hope for Depression Research Foundation. We thank Georg von Jonquieres and Matthias Klugmann from UNSW Sydney, Australia, for sharing the oligodendrocyte specific (MAG promoter) plasmids; Rachel Neve for the production of the HSV and Jill Gregory for her illustrations.

## Author contributions

OI and EJN designed the experiments and wrote the manuscript. OI, YYZ, DMW, ATB, CJB, RT, and JED performed mice surgery, behavioral studies, and tissue collections. OI performed the molecular analysis including RNA-sequencing library preparation and molecular cloning. AR and LS advised and performed bioinformatic analysis. SX, AKZ, and YD performed and analyzed the electrophysiological recordings. CT, WL, and CAT performed and analyzed the FISH study. MS supported IHC, confocal imaging, and quantification. BL generated the human RNA-seq dataset. JLD performed the EM myelin imagining and quantification. JWM provided the human blood samples.

## Competing interests

The authors declare no competing interests.

## Extended Data Figures legends

**Extended Data Fig 1.**
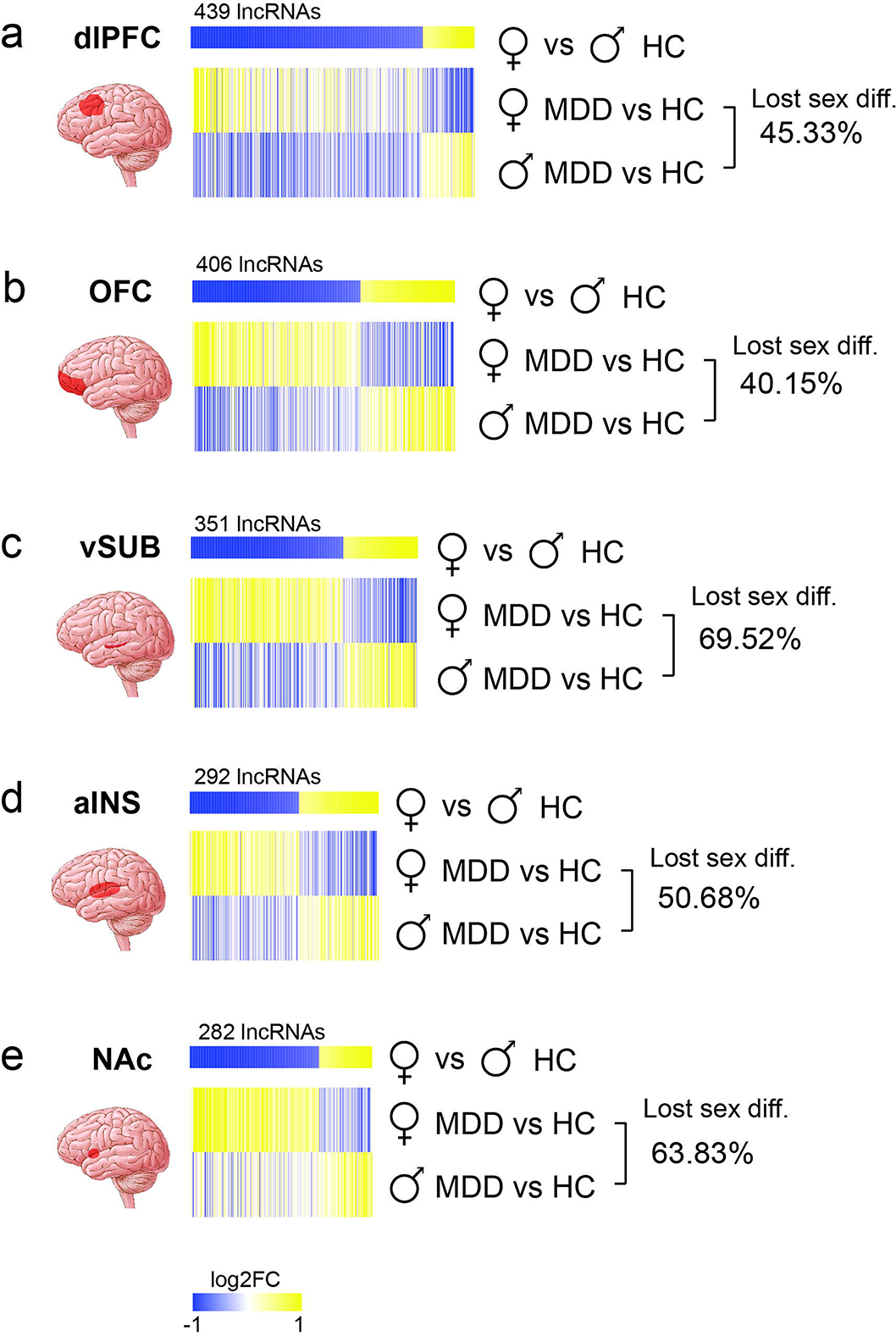
Loss of baseline sex differences in lncRNA expression in MDD. **(a-h)** Heatmaps demonstrating that lncRNAs that show baseline sex differences between female and male HCs (differential expression Fold change>30% and P< 0.05) are robustly oppositely altered in females and males with MDD compared to HCs. This pattern is noted in the in the dorsal lateral prefrontal cortex (dlPFC) **(a)**, orbitofrontal cortex (OFC) **(b)**, ventral subiculum (vSUB) **(c)**, anterior insula (aINS) **(d)**, and nucleus accumbens (NAc) **(e)**. The number of lncRNAs with baselines sex differences are indicated above the heatmap, as well as the percentage of lncRNAs that lose this sex difference in MDD. Yellow represents upregulation and blue downregulation between log2fold change=±1. RNA-seq dataset was originally published ^26^. n=9-13 per group. MDD = major depressive disorder; HC = healthy control.

**Extended Data Fig. 2.**
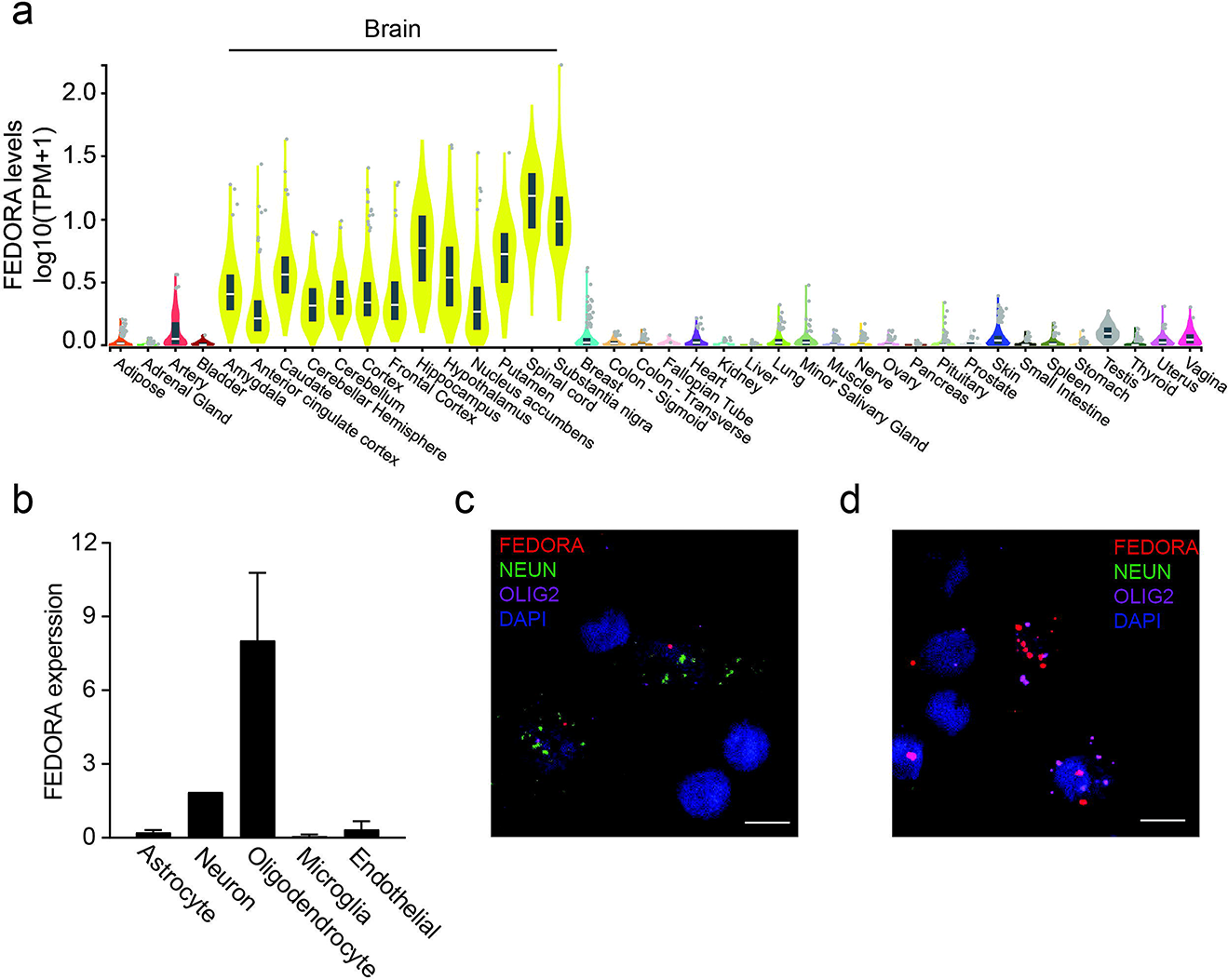
FEDORA is enriched in the human brain in oligodendrocytes and neurons. **(a)** Violin plots representing FEDORA levels across human tissue indicating brain enrichment. Data modified from ^22^. **(b)** FEDORA is expressed predominantly in oligodendrocytes as well as in neurons to a lesser extent. Data modified from ^23^. Bars represent mean ± SEM. **(c-d)** Images from fluorescent in situ hybridization on postmortem human brain rostral anterior cingulate cortex (rACC) tissue for FEDORA (red), neuronal marker (NEUN/RBFOX3 in green), oligodendrocyte marker (OLIG2 in purple) and cell nuclei (Dapi in blue). The images show nuclear localization of FEDORA and expression in neurons in gray matter **(c)** and higher levels in oligodendrocytes in white matter **(d)**. Scale bar=10µM.

**Extended Data Fig. 3.**
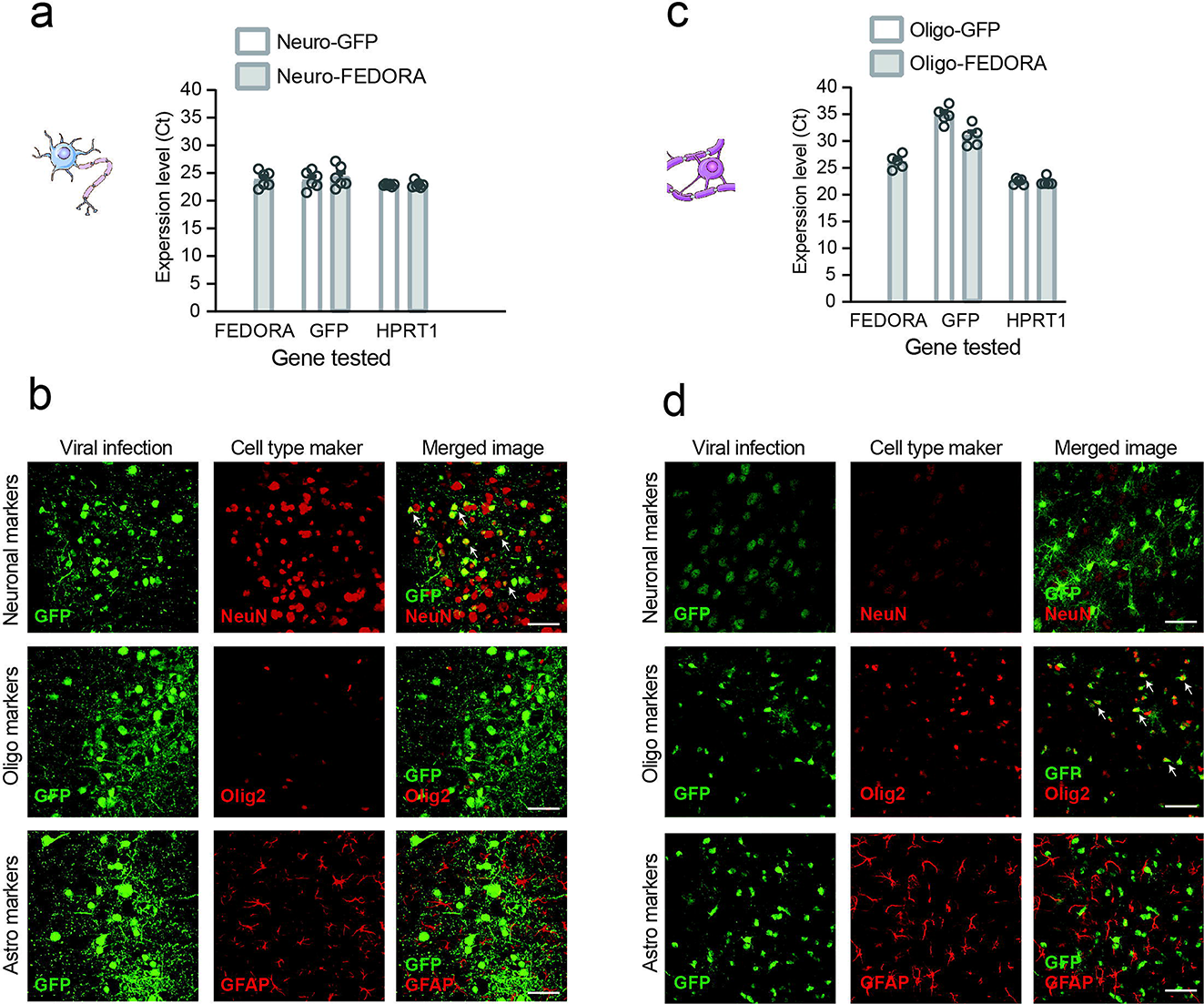
Validation of cell-type-specific FEDORA expression in mice mPFC. **(a,c)** qPCR results indicating Neuro-FEDORA **(a)** and Oligo-FEDORA **(c)** exclusively promote expression of FEDORA and similar levels of *Hprt1* and *Gfp* as the GFP control viruses. **(b,d)** Representative confocal images of immunohistochemical analysis of the cell-type specificity of the viral tools. Viral infection was reported in green labeling GFP (left columns) co-stained with red labeling of either NeuN for neurons, Olig2 for oligodendrocytes, or GFAP for astrocytes (middle column). The right column includes merged pictures indicating co-localized NEUN positive cells infected by Neuro-FEDORA **(b)** and Olig2 positive cells infected by Oligo-FEDORA **(d).** Examples of co-labeled cells are highlighted with a white arrow. Scale bar = 50 µM.

**Extended Data Fig. 4.**
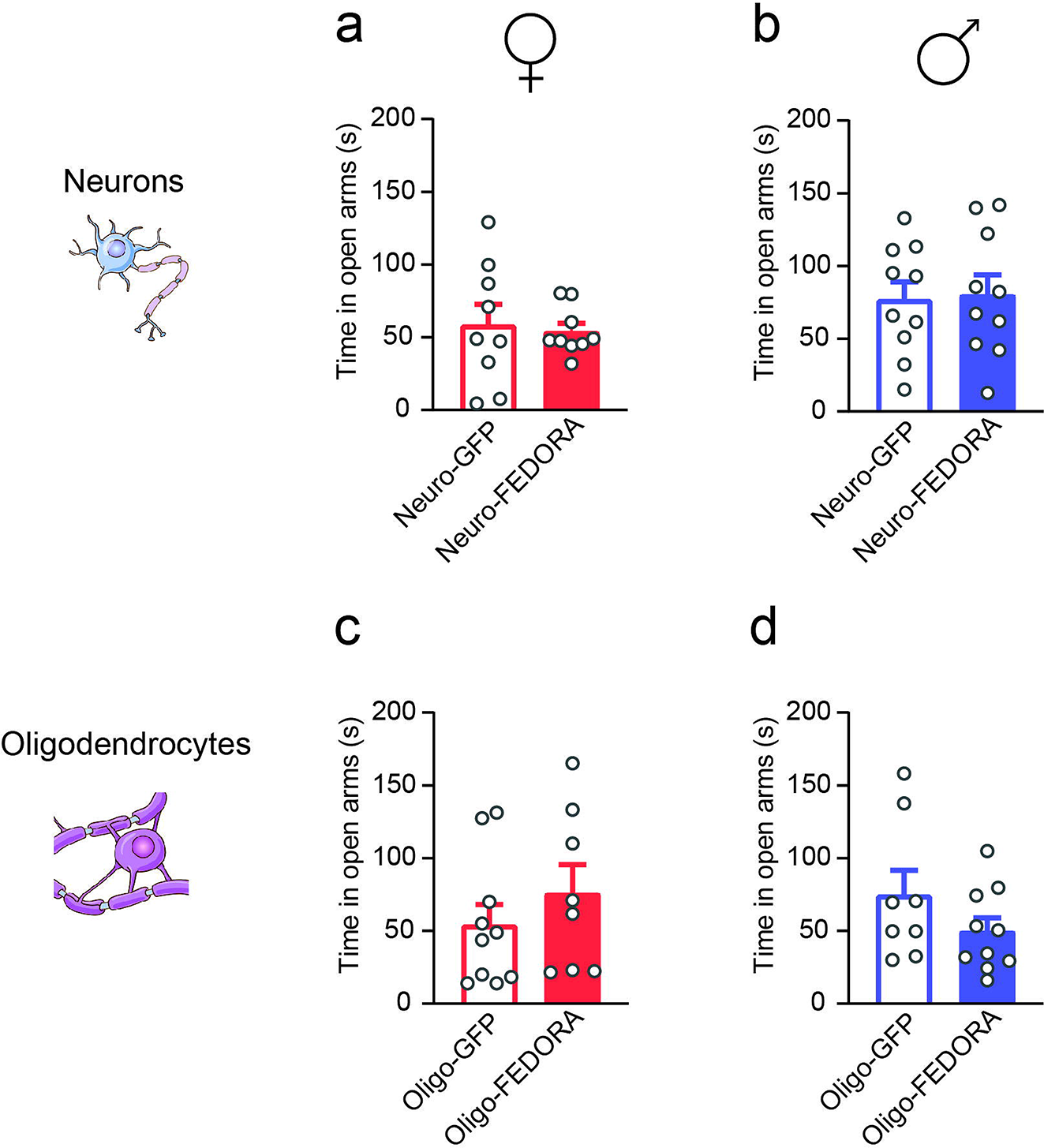
No effect of FEDORA expression in mouse mPFC in the elevated plus maze test. **(a,b)** Bar graphs showing that Neuro-FEDORA did not change the time spent in the open arms of the elevated plus maze compared to Neuro-GFP in either female **(a)** or male **(b)** mice. **(c-d)** Similarly, Oligo-FEDORA did not alter the time spent in the open arms of the maze in female **(c)** or male **(d)** mice compared to Oligo-GFP. Bars represent mean ± SEM and dots represent individual data points. N=8-10 per group.

**Extended Data Fig. 5.**
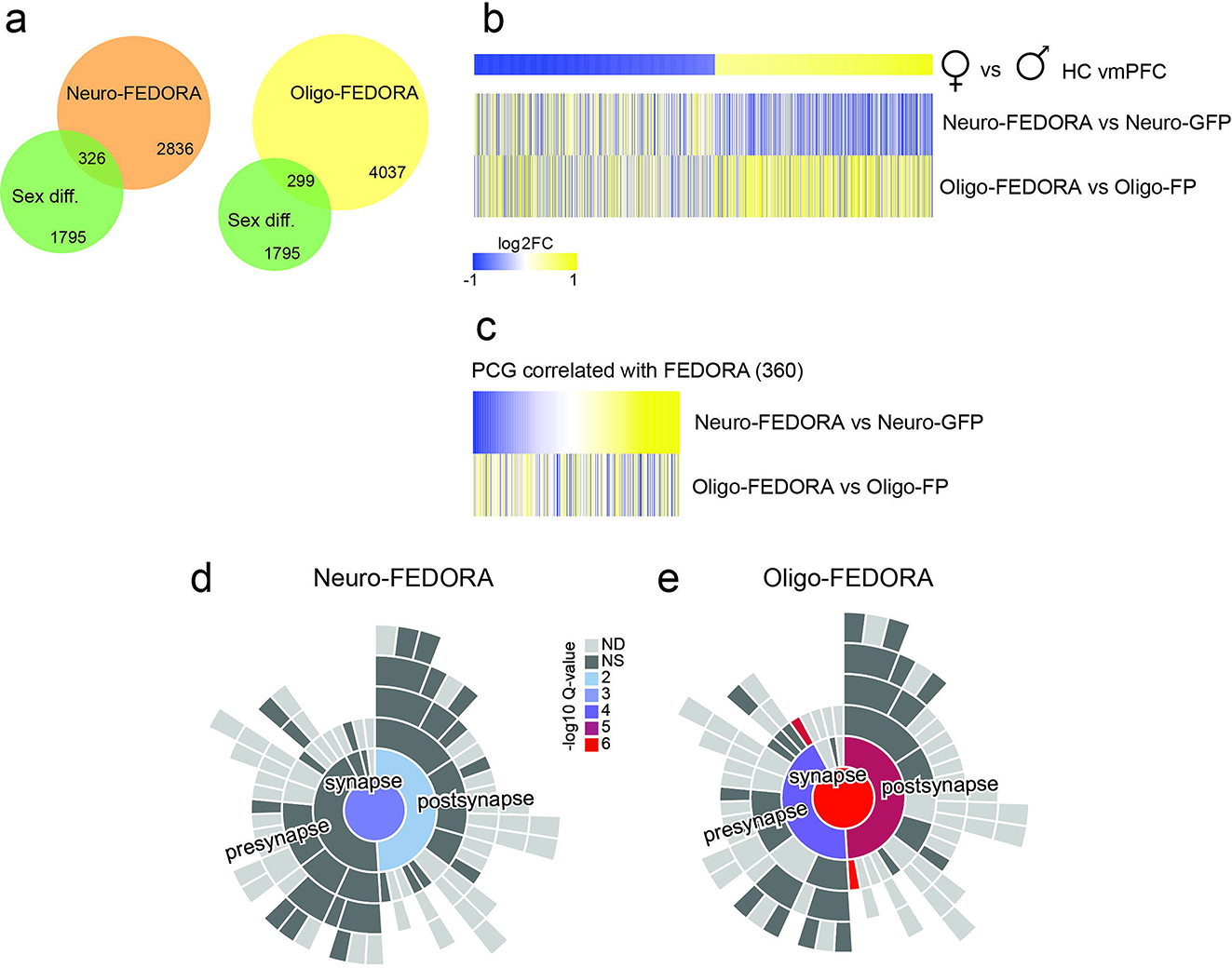
Cell-type-specific effects of FEDORA expression in female mouse mPFC. **(a)** Venn diagram indicating the overlap between genes regulated in female mouse mPFC by Neuro-FEDORA (left) or Oligo-FEDORA (right) vs mouse mPFC genes with baseline sex differences. **(b)** Heatmaps representing a loss of baseline sex differences upon FEDORA expression, mostly upon its neuronal expression. **(c)** Opposing regulation of the PCGs correlated with FEDORA in the human female brain by Neuro-FEDORA and Oligo-FEDORA. **(d-e)** Sunburst plot from SynGo ^91^ representing enrichment for regulation of synaptic and post-synaptic genes in Neuro-FEDORA **(d)** and synaptic, pre- and post-synaptic genes in Oligo-FEDORA **(e)**.

**Extended Data Table 1.** Demographics of the human postmortem subjects used for FISH study.

| **Subject** | **Sex** | **Age** | **Race** | **RIN** | **PMI** | **pH** | **AD** |
| --- | --- | --- | --- | --- | --- | --- | --- |
| CTRL1 | F | 15 | C | 8.9 | 25 | 6.3 | no |
| CTRL2 | F | 23 | C | 8.8 | 17 | 6.2 | no |
| CTRL3 | F | 45 | C | 8.4 | 17.3 | 5.9 | no |
| CTRL4 | F | 45 | C | 8.4 | 22.5 | 6.9 | no |
| CTRL5 | F | 50 | C | 6.2 | 23.3 | 6 | no |
| CTRL6 | F | 53 | L | 7.6 | 23 | 6.3 | no |
| CTRL7 | F | 60 | C | 7.9 | 20.3 | 6.4 | no |
| CTRL8 | F | 62 | C | 5.9 | 19.4 | 5.9 | no |
| CTRL9 | F | 68 | C | 8.8 | 16.6 | 6.4 | no |
| CTRL10 | F | 61 | C | 8.3 | 20.6 | 6 | no |
| CTRL1 | M | 13 | C | 8.4 | 25 | 6.7 | no |
| CTRL2 | M | 19 | C | 9 | 20 | 6.6 | no |
| CTRL3 | M | 20 | C | 8.2 | 21.2 | 6.8 | no |
| CTRL4 | M | 46 | C | 10 | 22 | 6.4 | no |
| CTRL5 | M | 48 | C | 10 | 14.7 | 6.8 | no |
| CTRL6 | M | 51 | C | 7.8 | 19 | 6.6 | no |
| CTRL7 | M | 54 | C | 10 | 19.3 | 6.7 | no |
| CTRL8 | M | 55 | C | 8.6 | 24 | 6.6 | no |
| CTRL9 | M | 59 | C | 8.7 | 22.2 | 6.4 | no |
| CTRL10 | M | 69 | C | 8.2 | 24.3 | 6.5 | no |
| MDD1 | F | 14 | C | 9.3 | 17 | 6.8 | no |
| MDD2 | F | 18 | C | 8.9 | 23.2 | 6.4 | no |
| MDD3 | F | 38 | C | 9 | 18 | 6.8 | no |
| MDD4 | F | 41 | C | 6.1 | 21.6 | 5.7 | no |
| MDD5 | F | 53 | C | 8 | 13.4 | 6 | yes |
| MDD6 | F | 58 | C | 6.3 | 14 | 6.4 | no |
| MDD7 | F | 61 | C | 8.9 | 22 | 6.7 | yes |
| MDD8 | F | 66 | C | 8.1 | 19.2 | 6.8 | yes |
| MDD9 | F | 66 | C | 4 | 17 | 5.4 | yes |
| MDD10 | F | 65 | L | 7.4 | 13.4 | 6 | no |
| MDD1 | M | 18 | C | 8.5 | 21.1 | 6.7 | no |
| MDD2 | M | 17 | C | 9.1 | 21.5 | 7 | yes |
| MDD3 | M | 20 | C | 7.6 | 17.3 | 6.7 | no |
| MDD4 | M | 44 | C | 9.1 | 19.5 | 6.1 | yes |
| MDD5 | M | 53 | C | 9.3 | 21.3 | 6.8 | yes |
| MDD6 | M | 53 | C | 8.1 | 24.5 | 6.4 | yes |
| MDD7 | M | 53 | C | 7.9 | 24 | 6.1 | no |
| MDD8 | M | 56 | C | 8.3 | 23 | 7 | yes |
| MDD9 | M | 56 | C | 7.4 | 18 | 6.3 | yes |
| MDD10 | M | 69 | C | 8.4 | 17.6 | 6.2 | no |

RIN, RNA integrity number; PMI, postmortem interval; AD, antidepressants.

**Extended Data Table 2.**
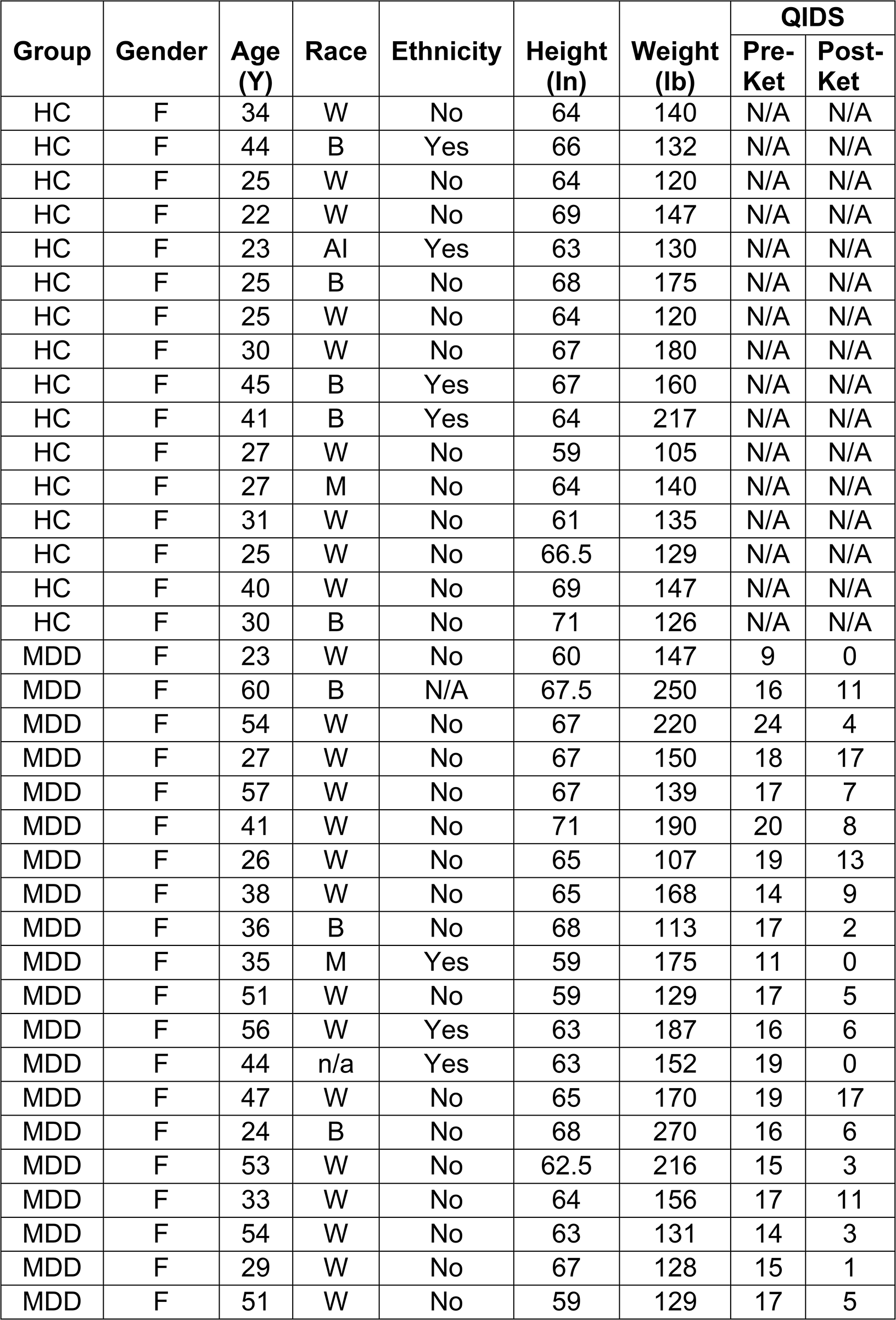
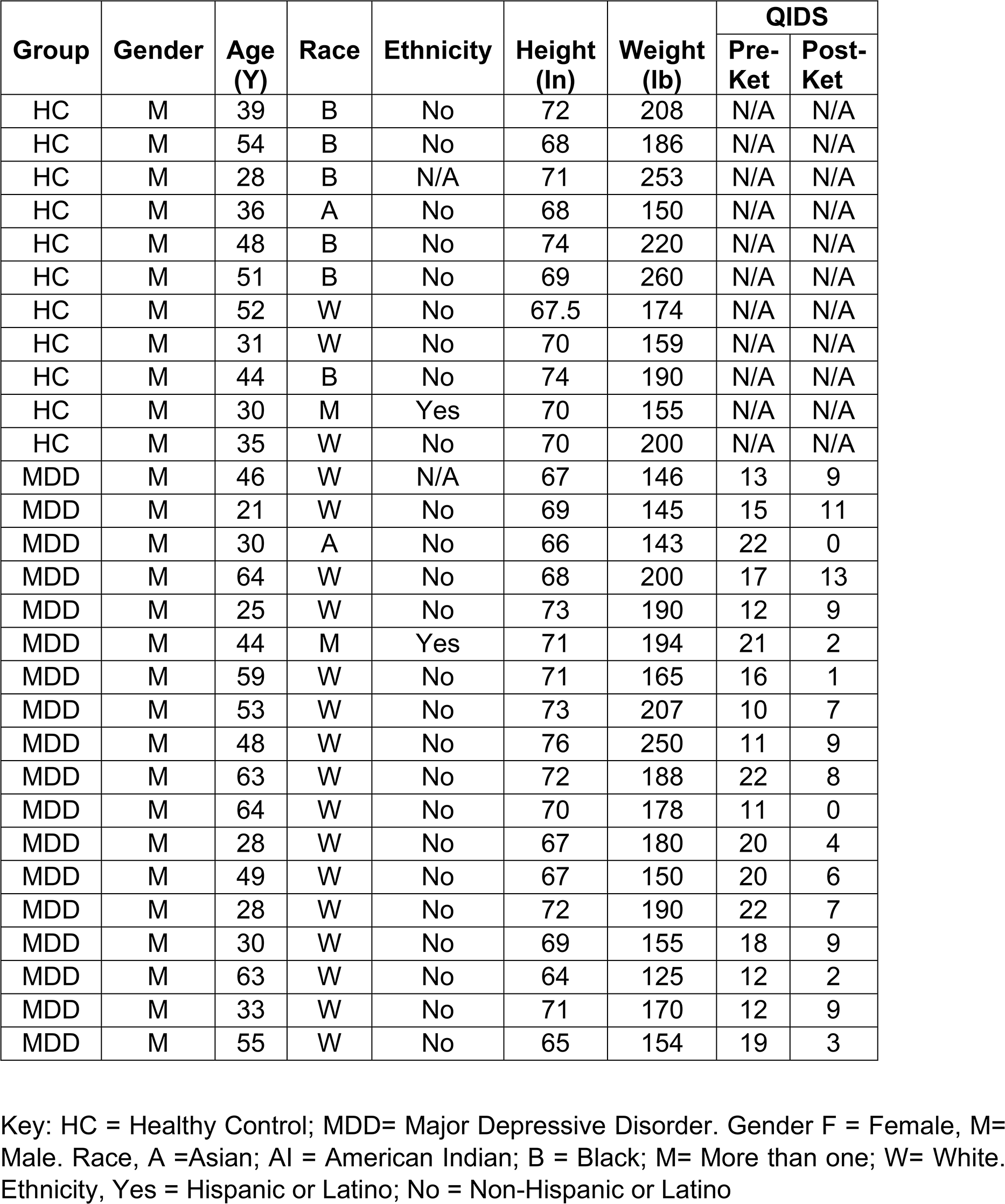
Demographics of the human postmortem subjects used for FEDORA blood analysis.

**Extended Data Table 3.**
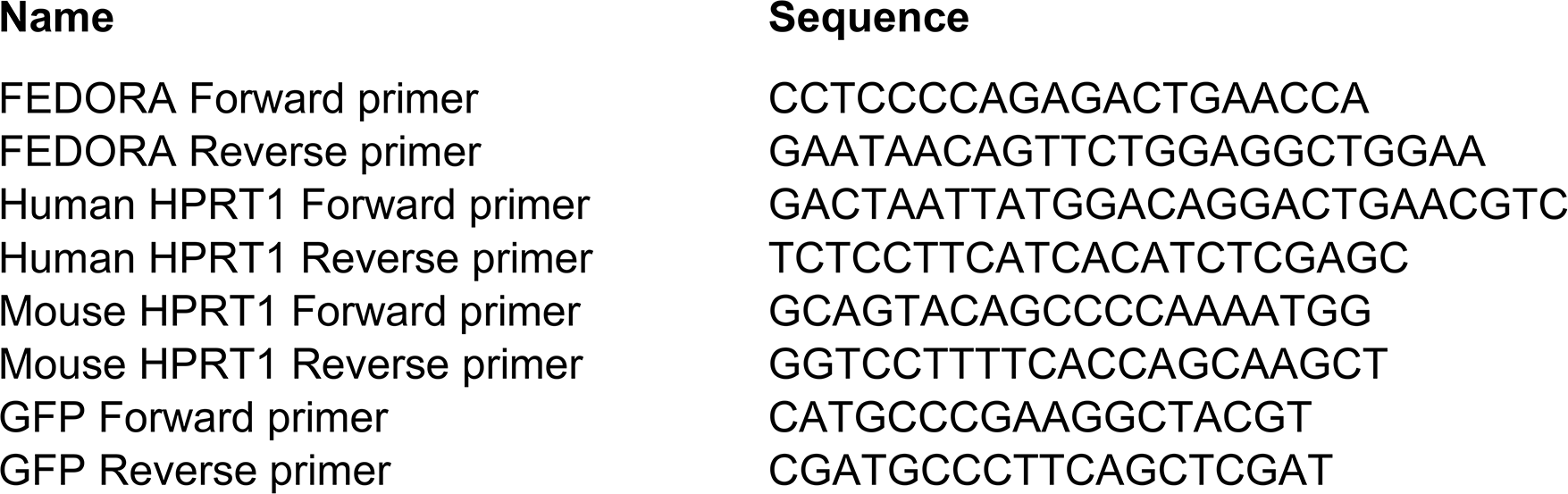
Sequences of qPCR Primers.

**Extended Data Table 4. Neuro-FEDORA vs Neuro-GFP differential gene expression list** (Enclosed excel sheet)

**Extended Data Table 5. Oligo-FEDORA vs Oligo-GFP differential gene expression list** (Enclosed excel sheet)

## Notes

### Competing Interest Statement

The authors have declared no competing interest.

